# A competitive advantage through fast dead matter elimination in confined cellular aggregates

**DOI:** 10.1101/2021.07.22.453340

**Authors:** Yoav G. Pollack, Philip Bittihn, Ramin Golestanian

## Abstract

Competition of different species or cell types for limited space is relevant in a variety of biological processes such as biofilm development, tissue morphogenesis and tumor growth. Predicting the outcome for non-adversarial competition of such growing active matter is non-trivial, as it depends on how processes like growth, proliferation and the degradation of cellular matter are regulated in confinement; regulation that happens even in the absence of competition to achieve the dynamic steady state known as homeostasis. Here, we show that passive by-products of the processes maintaining homeostasis can significantly alter fitness. Even for purely pressure-regulated growth and exclusively mechanical interactions, this enables cell types with lower homeostatic pressure to outcompete those with higher homeostatic pressure. We reveal that interfaces play a critical role in the competition: There, growing matter with a higher proportion of active cells can better exploit local growth opportunities that continuously arise as the active processes keep the system out of mechanical equilibrium. We elucidate this effect in a theoretical toy model and test it in an agent-based computational model that includes finite-time mechanical persistence of dead cells and thereby decouples the density of growing cells from the homeostatic pressure. Our results suggest that self-organization of cellular aggregates into active and passive matter can be decisive for competition outcomes and that optimizing the proportion of growing (active) cells can be as important to survival as sensitivity to mechanical cues.

## I. INTRODUCTION

Collective competition is ubiquitous in proliferating cellular systems, with implications ranging from maintenance of stable homeostasis in tissues [1], to robust morphogenesis [2], tumor development and suppression [3, 4] and survival of bacterial communities [5, 6], for example during range expansion [7]. Competitive advantages can be achieved either via direct adversarial interactions such as induction of elimination [8–10] (i.e. one cell inducing death in another) or due to competitive exclusion [11–15] where sharing a common resource or limited space leads to competition even without any specific response depending on the cell type of rival cells. The current work is aimed at the latter, non-adversarial, competition scenario.

While many past works focused on species’ typical division and death rates, arguing that fitness increases as the difference between the former and the latter [2], this intuitive argument cannot be used when cell aggregates do not expand but compete for space in a confined space, as validated numerically e.g. in Ref. [16]. In this case, division is, on average, balanced by death, creating a dynamic steady state known as homeostasis [4, 17, 18]. Then, the question arises whether an emergent property of the homeostatic state itself (in which competing cell types are absent) can instead serve as a suitable fitness indicator for competition outcomes. We ignore the role of chemical signalling, which can also participate in the competition that controls homeostasis [19]. In systems with pressure-regulated growth [20] and an equation of state relating pressure to the density of (living) cells, one property that has already been proposed is the homeostatic pressure [4, 16]. This makes intuitive sense, since, in a mixture of cell types with different homeostatic pressures, there is a range of intermediate pressures at which growth can still exceed death in one species while death exceeds growth in the other. However, an equation of state implicitly assumes that the active (growing) cells are the only mechanically significant component in the system, and thus the inclusion of any other continuously produced mechanical components such as dead cells or extracellular matrix may decouple the density of growing (active) cells from pressure and lead to a fitness that is not captured by homeostatic pressure.

Here, we demonstrate that the presence of passive by-products of the processes maintaining homeostasis can indeed significantly affect a species’ ability to compete. Specifically, we focus on systems where these by-products are the mechanical remnants of cells that have died, i.e. lost their ability to grow and divide, but are not removed immediately upon death. In real biological systems, the time scale for the removal of dead or dying cells can have a wide range of values relative to the cell’ life cycle depending on the mechanism of removal. Prominent examples are extrusion [17, 21, 22], structural disintegration [23], phagocytosis [9] or ‘explosive’ bacterial lysis [24]. Until now, models of growth-driven dense cellular matter have often implicitly employed instantaneous cell removal processes both in continuum theories and in most simulation works [25, 26] (and see also the discussion section). Taking into account this additional time scale is therefore not only theoretically interesting, but also of immediate biological relevance.

We consider a prototypical cellular system with only a few essential “building blocks” (Fig.1), namely steric repulsion, cell growth and proliferation, cell death, and for the sake of cell number homeostasis, also pressure regulation of growth [27–29]. The final ingredient of our model, which is central to our investigation, is the explicit, finite-time removal of dead cells, which translates to a continuous supply of (space-filling) dead passive matter and thus breaks the equation of state between pressure and the density of active, growing cells (Figs. 2A-2D).

**FIG. 1.**
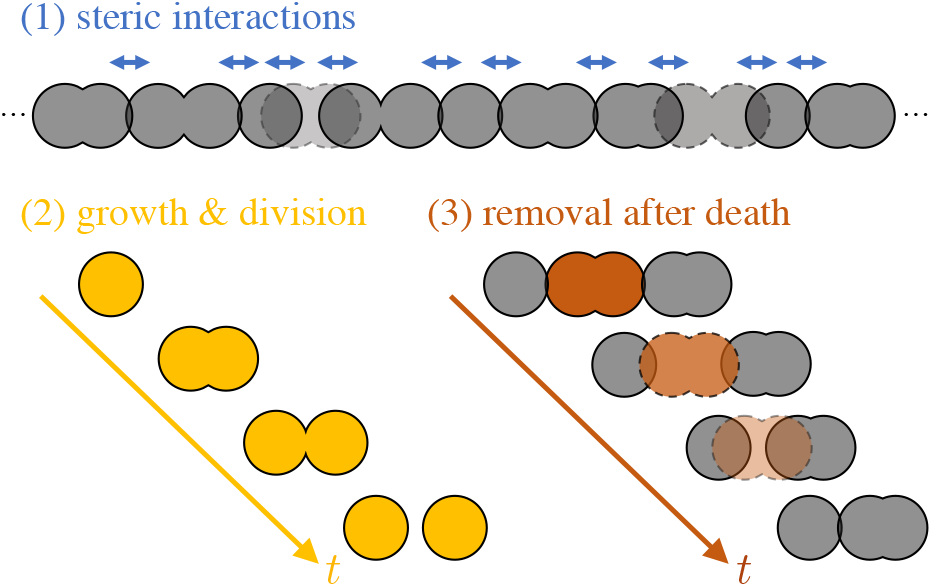
Essential “building blocks” for a cellular system with confinement induced-homeostasis. The presence of dead cells is explicitly taken into account as well. (1) Steric interactions push the cells to fill up the available volume (e.g. a 1D channel). Cells with dashed outlines are dead. (2) A schematic of cell growth and division for a single cell and it’s daughters. (3) Death and degradation schematic. Once a cell dies, its load bearing capability is progressively reduced resulting in a smaller effective cell volume. Not visualized: High density and pressure lead to inhibition of growth, thus regulating the balance between division and death which for a homogeneous sample results in homeostasis.

**FIG. 2.**
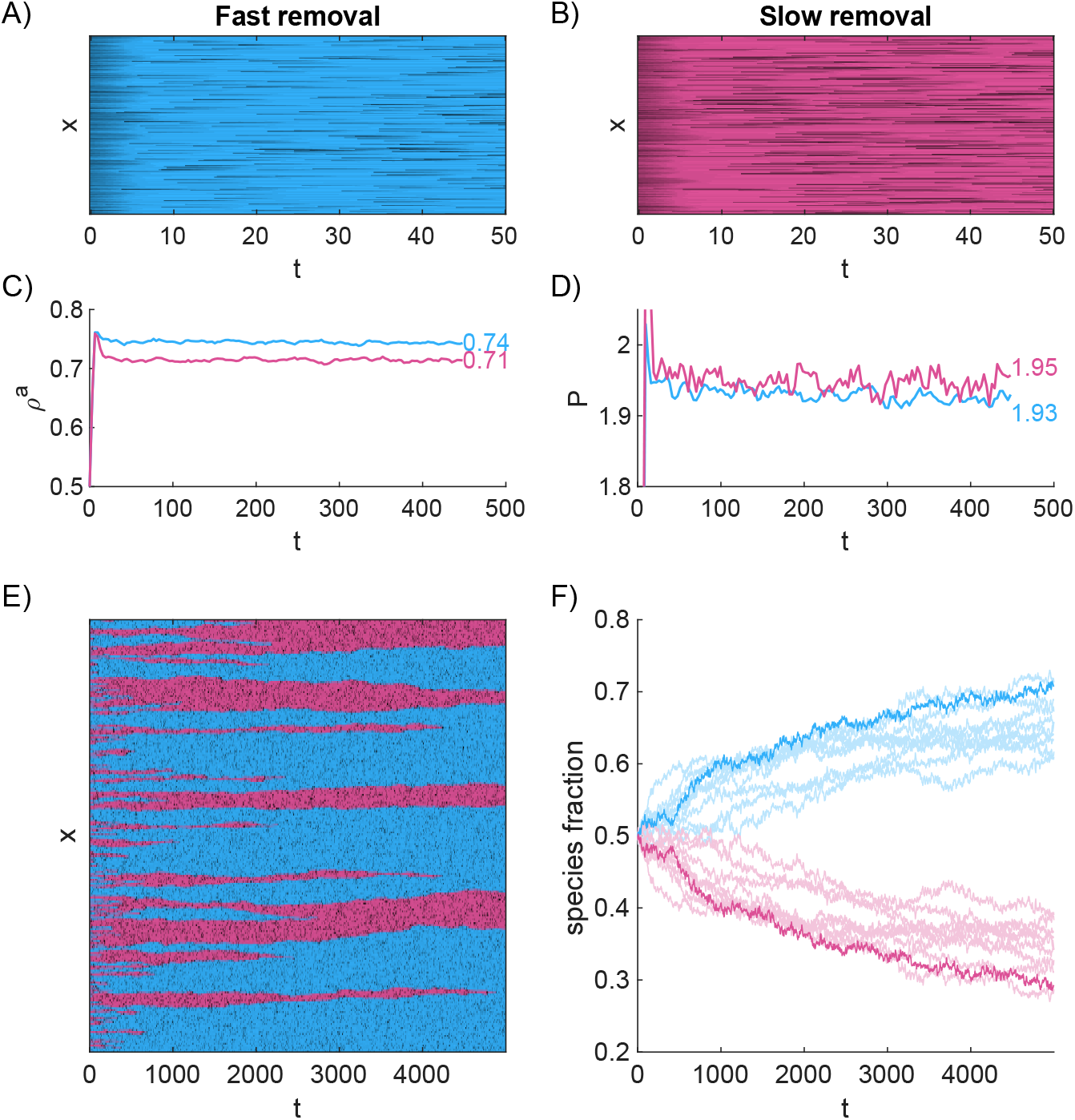
Representative numerical results for homogeneous samples and for competition in a mixture (Supplementary Videos 1 and 2). A,B) Coarse-grained kymographs for the living cells in homogeneous samples. H cells (blue) composing the system shown in panel A degrade more quickly after dying compared to L cells (magenta) composing the system in panel B. C) Time series of the density of growing cells (active density) for the homogeneous systems in panels A and B (averaged over 10 realizations for each). D) Time series of pressure for the same two systems (again averaged over 10 realizations). E) Kymograph for the living cells in a mixture of the two species starting at equal proportions (a subset of the volume is shown from a single realization). Spatial patches of species L are more susceptible to stochastic extinction. F) Time series of the population fraction in a mixture for 10 realizations of the same setup and parameters. The bolder line corresponds to the specific realization depicted in panel E. Species H consistently does better and takes over the sample.

**FIG. 3.**
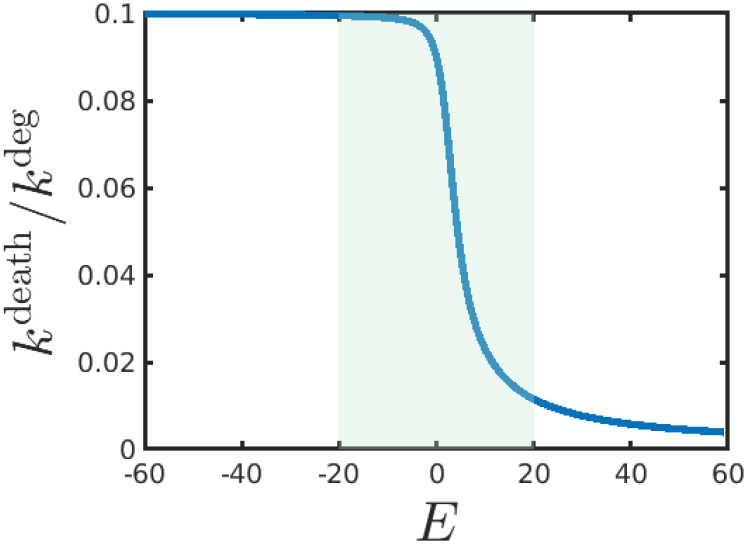
The inverse of the cell degradation rate *k*^deg^ as defined in Eq. (7) normalized by the death rate *k*^death^. The entire curve corresponds to the *E* range used in homeostasis simulations while the green shaded region denotes the range used for competition assays.

In such a system with by-products, we see that a species which removes dead matter faster and thus has a higher proportion of active, growing, cells can outcompete a species with a smaller active density even when mechanical pressure would suggest space invasion in the reverse direction (Figs. 2C-2F). This initial evidence raises the prospect of a complementary competition paradigm, which we will explore in this study using both theoretical considerations and systematic confirmation in simulations. We start with a simple mean-field continuum theory and show why this level of description does not reveal any particular advantage for species with faster dead matter degradation. However—keeping in mind that a well-mixed state, albeit an adequate description in a coarse-grained continuum model, is inevitably broken at the scale of finite cell granularity—we then provide a theoretical argument for how a fitness advantage could be induced by discrete cellular units, spatially heterogeneous growth rates and a finite mechanical screening length, and confirm that a Supplementary Video 1 for Fig. 2. Comparison of dynamics for two species (top: *E*_H_ = 4, 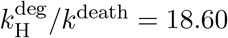; bottom: *E*_L_ = −20, 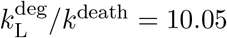) in separate single-species samples at homeostasis (a subsection of around 3% of the system length is shown for 200 time units). Dead degrading cells are colored black and denoted by a dashed outline and increased transparency denotes the current softening factor *S_i_*.

Supplementary Video 2 for Fig. 2. Dynamics in a competition assay between the same species as in Video 1 for 1000 time units at the very start of the simulation and displayed at x10 speed compatible spatial heterogeneity emerges in our simulations. Finally, we examine systematically in simulations how the active density is correlated to fitness and compare it to the fitness predicted by homeostatic pressure. We show that a homeostatic property, measured in a homogeneous cell sample, can determine fitness in competition and that competition is driven by interfaces between species.

## II. RESULTS

### A. Theoretical considerations

Before we present a detailed numerical analysis of our system, we first explore the consequences of passive matter production theoretically, in order to narrow down the possible mechanisms that can cause effects such as those seen in Fig. 2. The most general description of growing matter that adapts its growth/death dynamics to its local mechanical environment is on the level of continuity equations, which we first discuss in the context of a single species:

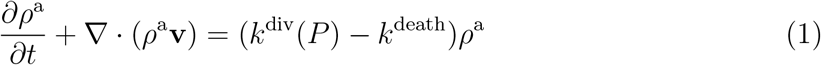

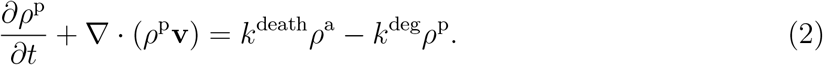

Here, the superscripts *a* and *p* indicate active cells (capable of division) and passive (dead) cells, respectively. We assume that all cellular matter has identical mechanical properties, where the isotropic local pressure is determined by an equation of state 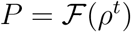 with the total density *ρ^t^* = *ρ*^a^ + *ρ*^p^ (for simplicity, we also assume that 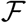 is an invertible function in the relevant range). In the over-damped limit, the velocity of the otherwise immotile matter is caused solely by intercellular forces according to force balance with a drag force *ζρ^t^***v** = –∇*P*, *ζ* being the drag coefficient. Therefore, both *P* and **v** are identical between Eqs. (1) and (2). Active cells constantly grow but reduce their growth rate in response to pressure, with a monotonically decreasing function for the growth rate *k*^div^(*P*). However, when cells die with rate *k*^death^, they do not vanish immediately from the system but are first converted to passive matter, which itself degrades with a rate *k*^deg^. These last two rates, that could in principle also depend on pressure, are taken as constant for simplicity. In a system of finite size, active density together with its passive remnants fills the system, growing with a local exponential rate *k*^div^(*P*) – *k*^death^, until growth is inhibited sufficiently by pressure.

Considering a mean-field approximation, where we assume that *ρ*^t^ and *P* are spatially uniform and thus **v** = 0, the steady state is determined by local homeostasis, which is defined by Eq. (1) as *k*^div^(*P*^h^) – *k*^death^ = 0, translating to a mean-field homeostatic total density of 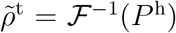. Meanwhile, Eq. (2) requires for homeostasis that 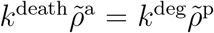, such that in steady state

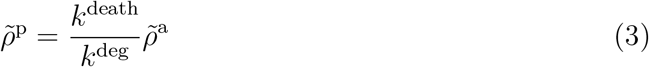

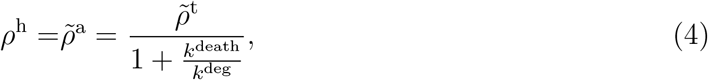

where we have defined the homeostatic active density *ρ*^h^ as a shorthand for the steady-state of the active density *ρ*^a^, as we will refer to it throughout this study. Note that, even if the mean-field quantities vary in time, 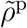 is still entirely determined by the history of the active density and approaches Eq. (3) on a time scale 1/*k*^deg^ if 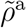 is constant. Therefore, we can view Eq. (4) also as a quasi-steady-state relation between *ρ^t^* and *ρ*^a^ when *ρ*^a^ changes very slowly.

We now turn to a competition scenario with two different species. From the mean-field consideration above, we see that *k*^deg^, the rate at which the passive matter is degraded, determines the relative proportions of active and passive density in homeostasis. In contrast, the homeostatic pressure *P*^h^ and the homeostatic total density 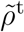 are solely determined by the condition *k*^div^(*P*^h^) – *k*^death^ = 0, independent of the amount of passive matter at this level of description. If we were to consider two species with different homeostatic pressures (e.g., through different *k*^div^(*P*)), a situation similar to that of Refs. [4, 16, 30] arises, where the species with higher homeostatic pressure has a clear growth advantage due to being able to exhibit net growth at higher pressures. In contrast, we consider here two species with dynamics as in Eqs. (1) and (2) that differ only in their respective passive matter degradation rates 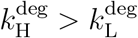, with 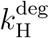 yielding a high active density (‘H’) and 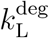 yielding a low active density (‘L’) in homeostasis according to Eq. (4) [note that an analysis in appendix F that goes beyond the noiseless continuum approach here predicts that different passive matter behavior can also induce slightly different homeostatic pressures due to second-order pressure fluctuations; however, as our numerical results below show, these are not the dominant drivers of competition].

Two species only differing in their passive matter degradation rates 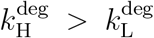 are characterized by two copies of equations (1), (2) for the densities 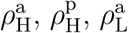 and 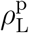. Again, **v** and P are both identical between all four equations. This leads to tight constraints on the possible relative behavior of the two species. For example, the mean-field version of Eq. (1), 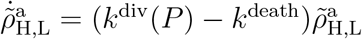, is identical for both the H and the L species, implying

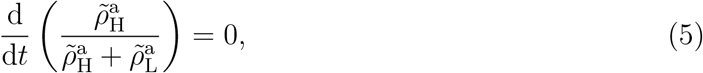

which excludes any meaningful competition, regardless of the behavior of the passive matter 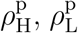. As long as the mean-field assumption of spatially uniform pressure applies, a similar equation holds for the total cell numbers, even if 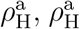 are spatially varying. By integrating Eqs. (1), (2), we have

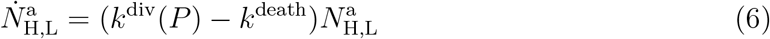

for both species, where 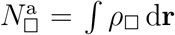 (with □ = *L* or H) indicates particle number corresponding to the relevant densities 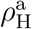 and 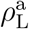, again implying the same relative change for both.

When the different species with their active and passive matter components are well-mixed on a microscopic scale, it is therefore not obvious how a species that leaves less passive matter behind could have a particular competitive advantage such as that seen in Fig. 2. Here, we argue that it is the granularity of the system at the cellular scale and its non-equilibrium nature that leads to a competitive advantage for the H species: Due to the growth and removal of discrete cells and finite mobility, the system never reaches a true steady-state, but is actively kept away from mechanical equilibrium (by which we mean the static configuration it would converge to if all growth/degradation processes where halted). It is easy to see that systems such as Eq. (1) have a well-defined length scale, or screening length, 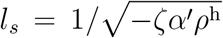 for pressure perturbations, where *α′* = d*k*^div^/d*P* is the growth response of the system at homeostasis (see appendix D). This screening length is the result of a competition between the diffusive spread of mechanical perturbations and the recovery of the cellular medium via adaptive growth/removal of matter which seeks to locally bring the system back to the homeostatic pressure *P*^h^. If *l_s_* is comparable to the cell size, a local pressure drop (for example, the removal of a cell) causes only a small number of discrete cells in its vicinity to grow in order to replenish the cellular material. At the interface between the two species, this finite-size selection can potentially lead to small-number effects. Indeed, a lattice-based thought experiment incorporating our most important assumptions—identical growth response for active cells of both species and different equilibrium fractions of passive matter—shows that there is a statistical bias towards finding fewer living cells of species L in this region than necessary to compensate cell removal (see appendix E), leading to a long-term takeover of species *H*. Therefore, species with a higher proportion of active, growing matter are statistically better at exploiting opportunities to ‘conquer’ new ground. This bias can also be shown to vanish for increasing screening length. Consequently, a finite screening length in conjunction with the discreteness of the system can yield a growth advantage for species H that is proportional to the difference in active densities 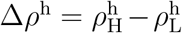.

We therefore expect that two species only differing in their passive matter degradation rate *k*^deg^ should have similar pressure-controlled growth rates during homeostasis when growing separately (because of identical homeostatic pressure *P*^h^). The same should be true for cells of both species when sharing space in the same system given they are far enough away from interfaces (because of a finite mechanical screening length). In contrast, based on the thought experiment mentioned above, growth rates near interfaces between the two species should be biased in favor of the species with a higher homeostatic active density *ρ*^h^.

### B. Numerical experiments

To test the correlation between homeostatic active density and fitness, we employ a minimal mathematical model which describes the dynamics of discrete cellular units and incorporates the above-mentioned building blocks (Fig. 1). In line with our theoretical considerations, we assume identical mechanical properties for all species and only a single differentiating feature: The degradation rate of passive cells. We then run single-species simulations to extract homeostatic properties and link them to the outcome of binary competition simulations.

#### 1. Agent-based numerical model

First, we briefly describe the cellular model used in the simulations and specifically the explicit cell removal process. The full details of the model can be found in appendix B. We describe a ‘dry’ cellular system where individual cells are modeled as immotile dumbbells, each composed of two nodes of radius R. Length is henceforth measured in units of node diameter. Nodes of different cells have a steric Hertzian [31] repulsion between them 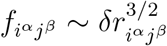 as a function of the node overlap *δr_i^α^j^β^_*. Here, Latin indices *i, j* denote cells, while Greek indices *α, β* distinguish between nodes of a cell. Nodes of the same cell have a non-linear spring interaction between them (also Hertzian), with rest length 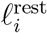. Cell growth is implemented via elongation of the internal spring’s rest length 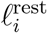, expressed equivalently via a growth phase 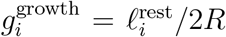 in the range [0, 1]. A cell starts its life with two nodes fully overlapping 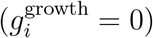 and, if it does not die before then, ends it with division when 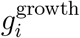 reaches 1 and the two nodes just touching (zero overlap), as depicted in Fig. 1.

Each cell has a base growth rate 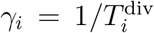, where 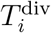 corresponds to the division time in the absence of external forces. We choose this basal growth to be significantly faster than the death rate *k*^death^ (see further below) which cells will have to approximately match in homeostasis by decreasing their growth rate. For this purpose, the base growth rate is modulated by a monotonously decreasing sigmoidal function of the cell-sensed pressure 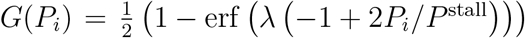 (see appendix B for details), making the overall growth rate of 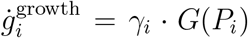 dependent on external forces and thereby allowing for homeostasis. *λ* determines how steeply *G*(*P_i_*) decays and *P*^stal1^ is the pressure at which cell growth effectively halts. A visualization of *G*(*P_i_*) is provided in appendix B. Upon division, each node of a mother cell turns into a new daughter cell 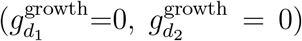, itself composed of two connected fully overlapping nodes. To avoid synchronization of divisions due to lineage extinctions, the new base growth rates *γ*_*d*_1__, *γ*_*d*_2__ are randomly drawn from a normal distribution (as shown e.g. in bacteria in Refs. [32, 33]). Death of a cell is stochastically initiated for each cell at a rate *k*^death^ that is much lower than the average base growth rate and the cell degradation rate (see below). The activity of the cells, and any stochasticity, is present only in their growth, and their division and death events. Growth and division, coupled to the inter-cell steric repulsion, result in local forces which can be relaxed to a certain extent through cell motion (see over-damped equations of motion in appendix B) but which, in a closed system, lead to an overall pressure buildup. This in turn feeds back on growth which leads to a dynamic state of homeostatic balance between cell division and death.

To the quite standard model described so far, we now add a new mechanism of mechanical cell degradation and removal that has an explicit and tunable time scale, via node “softening” (see Fig. 1). For this purpose, each cell is assigned a factor *S_i_* which scales the default Hertzian inter-cell forces: 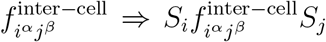 (see Eq. (B6) in appendix B). For a living growing cell, *S_i_* = 1. However, once death is initiated, growth stops irreversibly so that the cell turns from active to passive matter. The scaling factor *S_i_* then decays according to 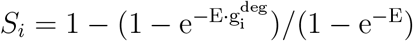 (see visualization in Fig. 8) with progress tracked by a degradation phase 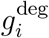 that grows linearly in time from 0 to 1. The parameter *E* is used to control the softening profile and is shown below to control the rate of the mechanical degradation of the cell *k*^deg^. After a fixed duration *T*^removal^, also taken to be significantly shorter than 1/*k*^death^, *S_i_* fully decays to zero at which point the cell is completely removed from the system.

While the above numerical model can be easily generalized to 2D and 3D, here, the long-axis orientation of the cells, their growth and their motion are all constrained to a continuous 1D periodic domain (see Fig. 1). We use a channel length of size 2560 in node diameter units, which is several orders of magnitude larger than the screening length (cf. appendix D), eliminating finite size effects, and also large enough to provide sufficient statistics (cf. appendix. The initial seeding is always done with a number density of 0.5.

To capture the shortest time scale in our system—the degradation time of cells—we chose *T*^removal^ = 10, such that cell degradation happens on the order of our model time units. The generation time in our model is determined by the death rate, since the growth rate of cells *γ_i_* · *G*(*P_i_*) decreases to approximately match it during homeostasis. For the degradation time to be significant, but substantially shorter than the typical doubling time, we fixed *k*^death^ at 10^-2^, leading to an approximate doubling time of ln(2)/*k*^death^ ≈ 70 or 14 doublings per 1000 time units (see appendix F for details on the expected homeostatic pressure underlying this resulting growth rate). The full set of parameters can be found in appendix B.

### C. Single species homeostatic properties

To measure homeostatic properties such as *ρ*^h^ and *P*^h^ for a given cell species, we run simulations of the above model, each with a single species, and as before, the species differ only in their softening factor *E*. Each such simulation is run for a time 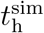 of 2000, corresponding to approximately 30 generations. Time-averaged measurements, such as for the homeostatic properties *ρ*^h^, *P*^h^, are performed in a time window 1000 < *t* < 2000, well after the steady-state is reached (see Fig. 2). In the following, we use the brackets 〈 〉, to denote an average over an ensemble of realizations. As a measure for the cell degradation rate we chose the time it takes the force scaling factor *S_i_* to drop by 90% after death:

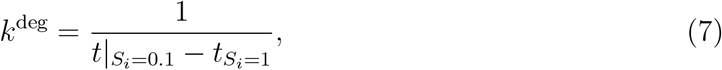

representing the fact that despite the fixed *T*^removal^, cells are effectively removed already when their *S_i_* value decays close enough to zero.

An intuition for the general behavior of our system can already be gained from the representative examples shown in Figs. 2A-2D (Supplementary Video 1) in light of the theoretical considerations presented above. Figs. 2A and 2B visualize coarse-grained spatio-temporal dynamics for two separate cell populations by tracking cell positions through time (only the first 50 time units out of 2000 are shown). Colored regions represent living cells and any black spaces in between denote either voids or degrading (dead) cells. Fig. 2A corresponds to a species H with *E* = 4 and thus a faster degradation rate 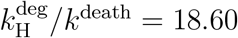 compared to *E* = −;20 and 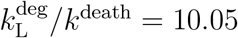 for species L in Fig. 2B. The seeded cells fill up the available volume already within 5 time units. The higher density of black lines that is apparent for species L in the steady state represents a greater accumulation of passive matter and supports our expectation from Eq. (4) that the homeostatic active density for species L is lower than for species H. Indeed, the time series for the active density in Fig. 2C, measured as the number of growing cells divided by system length (averaged over 10 realizations, only the first 500 time units are shown), clearly confirms this as does the homeostatic value (denoted in the figure).

It is also useful to consider the pressure, since it is the mechanical property that controls growth. While, a priori, we do not expect an inherent difference in homeostatic pressure (see theoretical considerations above), the time series in Fig. 2D indeed shows a small but significant difference between the two species. The pressure is measured via the Irving–Kirkwood expression [34] [35] which in 1D takes the form: 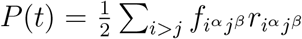 (averaged over 10 realizations in the figure) with *α* and *β* denoting nearest nodes of the neighbor cells. The homeostatic value *P*^h^ in this specific case is slightly higher for species L compared to H as denoted in the figure.

We next measure these two homeostatic properties for species with a range of *E* values and associated degradation rates. One can immediately see in Fig. 4 that they show systematically different behavior across the parameter space: The homeostatic active density shows a monotonous increase with *E* and is predicted exceptionally well by the mean-field expression in Eq. (4) [36]. The homeostatic pressure, on the other hand, is not constant as the mean-field consideration predicts (although here it increases by only ~ 2% of its minimal value). The reason for this are *local* pressure fluctuations generated by degrading dead cells. This is most evident when considering the *E* → ±∞ limits where a cell vanishes instantaneously and the pressure drops abruptly. Here, the measurements performed for *E* < −30 and *E* > 30 correspond to a regime in which the mechanical collapse of the cell is indeed much faster than the recovery response of the surrounding cell medium (*S_i_* approaches a step function; see appendix B). One should note that the fast cell collapse limit is not a peculiarity of the current model but in fact common practice in studies of competition and cellular active matter in general which use it implicitly by removing cells instantaneously [4, 16]. At the other extreme of the spectrum, the pressure should be most closely approximated by the mean-field prediction when the local pressure fluctuations are negligible (compared to the mean value). For the numerical model employed here, the *global* pressure fluctuations are minimal at *E* ≃ 4 (see appendix F), where indeed also the pressure is minimal: min (*P*^h^) = 1.924, and away from this point increased fluctuations lead to increased homeostatic pressure. The zeroth order mean-field value for the homeostatic pressure is 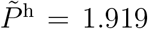 (dashed red line in Fig. 4B; see appendix F for derivation) which is even lower than the minimum measured in simulation, compatible with the fact that the local pressure fluctuations are never truly negligible in this numerical model. This observed deviation of *P*^h^ from the zeroth order mean-field calculation is in fact useful and will allow us to extract an alternative fitness prediction from the homeostatic pressure [4, 16] that can be compared to that of the active density.

**FIG. 4.**
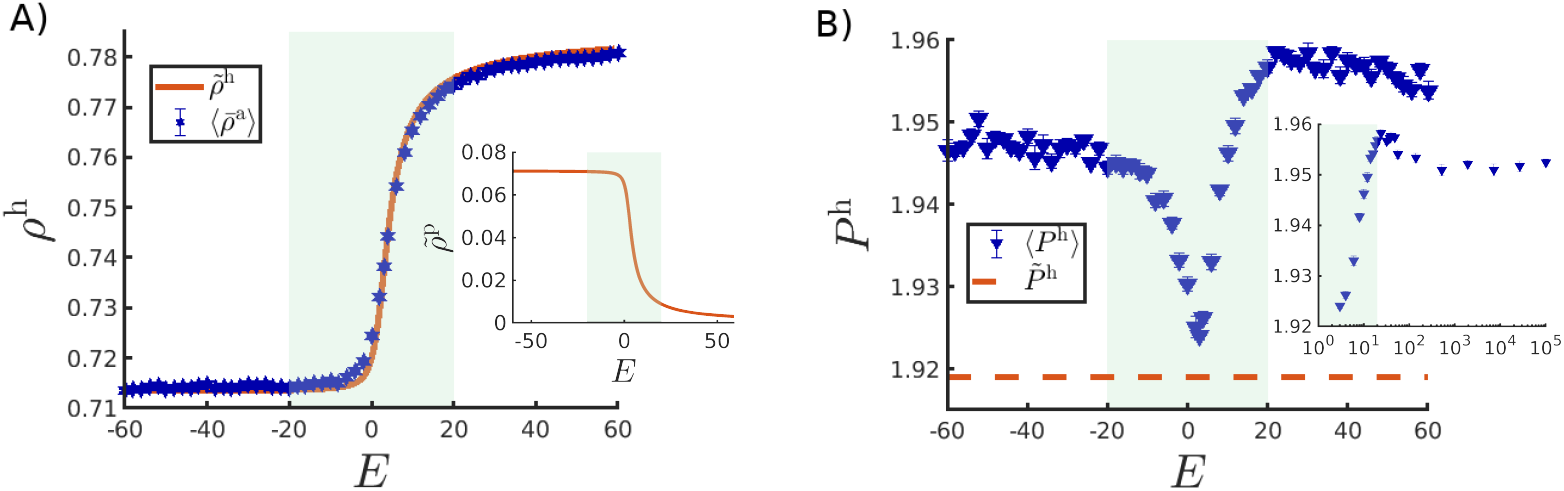
Possible competition drivers as a function of the softening factor *E* measured in homeostasis (averaged over 10 realizations). The green shaded regions denote the range of *E* values used in the competition assay of Figs. 5 and 6. A) The homeostatic active density (blue stars, error bars are smaller than marker size). The orange line is the prediction of Eq. (4) using a reference total density of 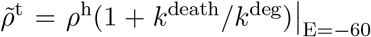. Inset: Mean field prediction for the passive density based on Eq. (3) and the 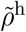 shown in the main figure. B) Homeostatic pressure (error bars are smaller than marker size). The dashed orange line is the mean-field prediction 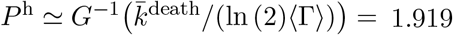 (see appendix F). Inset: same in log scale for *E* ≥ 3.

### D. Comparison of competition drivers for cell mixtures

To determine how these homeostatic properties are correlated with species fitness, we perform pairwise competition assays between the species differing in their softening factors *E* and associated passive matter degradation rates, and observe the relative change in species abundance over time. In each simulation, the channel is initially seeded with a balanced mixture of both species in a random arrangement.

A representative example of such a competition assay, which pits a species L against a species H with a higher degradation rate, is shown in Figs. 2E-2F (Supplementary Video 2). From the spatio-temporal dynamics of the cells shown (coarse-grained) in Fig. 2E we see that, besides general coarsening dynamics, patches of both species H and L grow and shrink stochastically. Yet, species H has a clear statistical advantage, as quantified by the species fractions *N*_H_(*t*)/*N*(*t*) and *N*_L_(*t*)/*N*(*t*) in Fig. 2F. Longer simulation runs show that such winner species eventually take over the entire sample (see appendix G). The observation that a species H with a higher homeostatic active density wins the competition is compatible with the theoretical considerations presented above.

In the competition assays for other *E* value combinations, we continue to label cells with a higher *E* value as species H and those with lower *E* as species L. We focus on the parameter regime in which we expect the clearest distinction between the two predictions; namely −20 ≤ *E* ≤ 20 (denoted by green shading in Fig. 4) where both homeostatic pressure and active density change significantly and where the functional form of *S_i_* is more sensitive to changes in *E* (see appendix B). For reference, we also run simulations where all cells have the same *E* value and arbitrarily label two equal size subsets of the seeded cells as H and L. Measuring “fitness” for such fully neutral competition allows us to estimate the natural statistical variability that arises from the model and from the chosen protocol of fitness measurement. We define relative fitness for species H as the fraction of the final proportion of cells 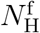 in the entire population *N*^f^, measured after a fixed time 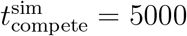 (~ 70 generations). This simulation time was selected to be long enough to show a clear trend for the change of species fraction (cf. Fig. 2F), but not long enough to reach the absorbing extinction boundary, which would have biased the measurement [37]. The relative fitness measure 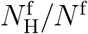 averaged over an ensemble of simulation realizations is shown in Figs. 5A-5B. As expected, the species fraction in neutral competition (equal *E* values) is 0.502 (±0.005). At the other extreme, the relative fitness in this numerical experiment grows up to a maximum of 0.75 (±0.02) showing that some species carry a distinct advantage over others.

**FIG. 5.**
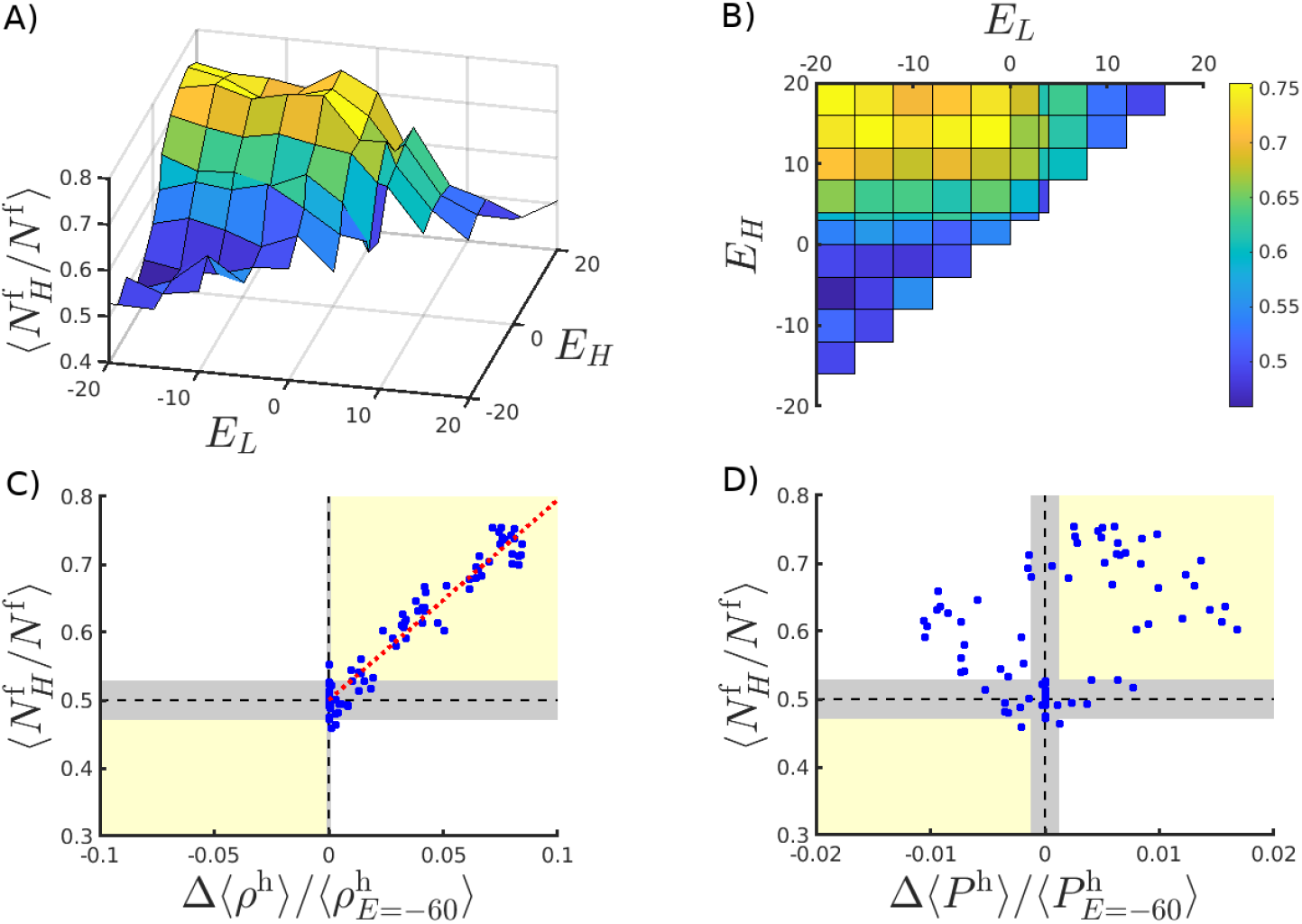
Competition and fitness. A) Relative fitness (species fraction at the end of a competition assay) of two species H and L differing only by their *E* values. Height and color (see color bar in panel B) both indicate the fraction of cells of species H out of the total population after 5000 time units. B) Color-coded 2D top view of the same plot. Fitness in competition vs fitness predictions: C) Fraction of species H (at end of a competition assay) vs the difference in active density between the corresponding single-species homeostasis assays 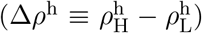, normalized by the value of *ρ*^h^ measured for *E* = −60. The dotted red line is a linear fit with a slope of 2.95 and a root mean square error of 0.02. D) Same but with respect to the difference in homeostatic pressure between species. In panels C,D, the yellow colored quadrants (delimited also by black dashed lines) indicate a successful prediction of competition (see text) and the grey areas delimiting the quadrants provide a estimate of the statistical variability. For the *y* axis, the variability estimate is defined by the range of *all* data points at *E*_H_ = *E*_L_ (where the correct *y* value is trivially 0.5). For the *x* axis, we define the variability margins as twice the standard deviation of the relevant property (〈*ρ*^h^(*E*)〉 or (〈*P*^h^(*E*)〉 in the regime −60 ≤ *E* ≤ −30 (where they level off, see Fig. 4). Data points in all panels correspond to an average over 9-10 realizations each.

Across the parameter space, the species labeled H exhibits either a similar or a higher fitness than species L, indicated by 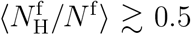 ( Figs. 5A-5B). It is also worth noting that, in the range −20 ≤ *E* ≤ −5, there is no clear advantage for any of the species, although the homeostatic pressure changes considerably in this range (Fig. 4). These two observations already indicate a dominant role of the homeostatic active density.

For a more quantitative analysis, we correlated the final cell fraction with the difference in homeostatic active density between the competing species. We do the same for the homeostatic pressure and in both cases normalize by a reference value at *E* = −60. The results of these two measurements are shown in Figs. 5C-5D. For the active density (Fig. 5C), all the data points fall in the quadrants of positive correlation (yellow shading) within statistical variability. A clear linear trend indicates that the fitness as defined by the species fraction can accurately be predicted from the difference in homeostatic density. From a different perspective, a larger homeostatic active density difference leads to a faster average rate at which the winner species takes over the sample, in line with our theoretical framework. The linear dependence can be fitted as

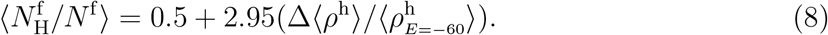

The root mean square error of the fit is 0.02 which matches the scatter of data points for identical species (i.e. at Δ*ρ*^h^ = 0). For the homeostatic pressure (Figs. 5D), an equally unambiguous correlation is not observed, even taking into account statistical variability (greyed out areas in the figure). For certain mixtures, a species with a lower homeostatic pressure can in fact win the competition (upper left quadrant), as already seen in Fig. 2. We thus conclude that, in this competition scenario, the homeostatic active density has a higher impact on fitness than does the homeostatic pressure.

Finally, we examine another prediction of our theoretical consideration, namely that preferential growth of a winner species happens only near interfaces between species. For this purpose, we define ‘interface’ cells as ones that are less than a distance 15 from a cell of another species (either active or passive) and measure their average growth rates starting from a time of 1000 (before which the system has not coarsened enough to provide good statistics for ‘bulk’ cells). Figs. 6A-6B shows that growth rates at interfaces increase with the difference in *ρ*^h^ for species H and decrease for species L. In contrast, ‘bulk’ growth rates do not show any strong trend. A typical time trace of the different growth rate averages for a case of high homeostatic active density difference is shown in Fig. 6C, indicating that this advantage, despite considerable stochasticity and the general coarsening dynamics (cf. Fig. 2), is present throughout the simulation. Collectively, this evidence suggests that preferential growth and consequently the competitive advantage are indeed induced by interfaces, supporting our proposed ‘opportunistic growth’ mechanism.

**FIG. 6.**
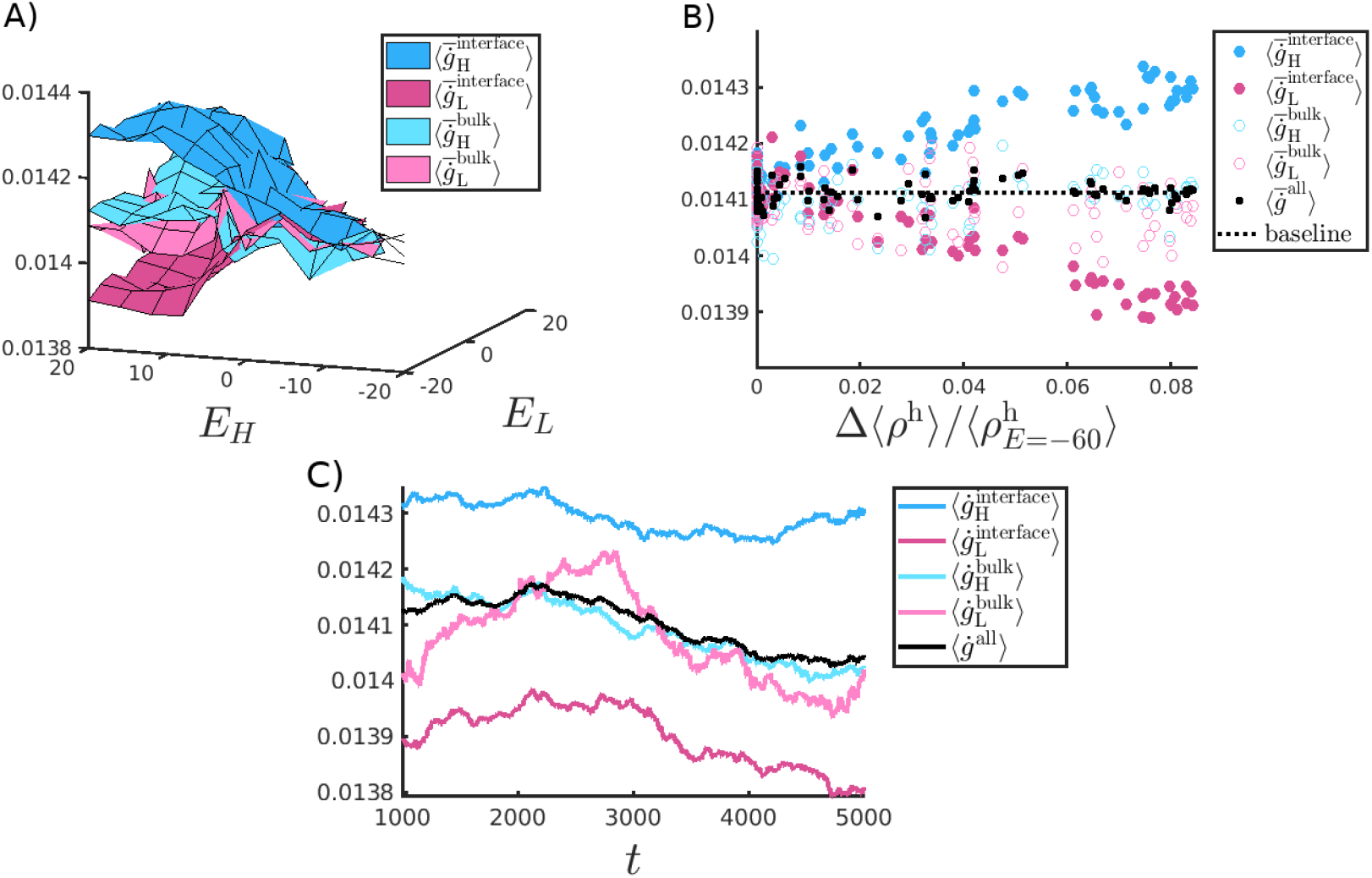
Growth at interfaces. A) Growth rates of competing cells near and away from species interfaces (with axes *E*_A_, *E*_B_ rotated compared to panels A,B of Fig. 5 for clearer visualization). B) Same growth rates plotted vs the (normalized) species difference in active density. The black dots denote the average growth rate of all cells in a given sample (averaged over realizations), and the dotted line denotes the mean of this quantity for all species pairs. Clearly the preferential growth through which a species H manages to overtake its competitor L happens at the interfaces between the two species. C) Time series of the different growth rates for two specific species *E*_L_ = −20 and *E*_H_ = 16 that have a high difference in their homeostatic active density 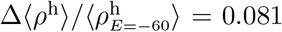. Growth rates in panels A,B are averaged over the time range 1000-5000 and in all panels over 9-10 realizations.

## III. DISCUSSION

In this work, we studied collective competition in growing cellular matter; a common phenomenon in living systems that is responsible both for helpful processes in an organism, such as maintaining homeostasis in an organ, as well as in harmful ones, such as tumor development. We addressed a type of competition between cell types that results from purely mechanically regulated growth, independent of the cell type of interaction partners, i.e. without any adversarial response aimed specifically at competing cell types. Instead, we considered a scenario of competitive exclusion where cells in confinement vie for room to grow. We found that the fitness of a given cell type, which indicates how it will fare during competition, can be determined from its emergent homeostatic properties in a homogeneous system where no competition takes place. In our case of pressure-regulated growth and different passive matter degradation rates, the decisive property turned out to be the density of growing cells (active density).

Evidence for homeostatic cell density as a predictor of fitness can be found in both experiments and simulations [1, 16, 38, 39] and indeed homeostatic density has already been suggested as a competition driver, but only in a very limited sense: one that attributes fitness to a species-dependent threshold density above which apoptosis is triggered [40–42]. The quite intuitive reasoning for the homeostatic density driving the competition in that specific case is that the “loser” species can be above the threshold and diminishes while the “winner” species is still below its own apoptosis threshold and prospers. Here, on the other hand, we allow for a more general relationship between the death and division rates that still supports homeostasis. Another distinction is that Refs. [40–42] considered division and death rates to depend directly on the density of growing cells, while, here, we consider these to depend on pressure as seen experimentally in many systems such as epithelia [29], tumours [43, 44], morphogenesis [2], yeast [28] and bacteria [45]. When proliferation and/or removal rates depend on pressure, they are affected by the production of all load-bearing matter in the confined system, and not just by the active growing cells. Thus, the continuous production of passive matter such as dead cells or extracellular matrix, as well as its removal process, can affect the cells’ ability to compete.

Here, specifically, we considered the case of dead cells, by replacing the instantaneous removal employed in many other works with a finite-time degradation process in both theory and an agent-based numeric model. This has the effect of introducing an explicit, tunable time scale for the persistence of passive dead cells that is separate from the time scale at which cells stop growing and die, and which results in a finite density of passive dead matter in the system.

We showed that a mean-field consideration in the continuum model—describing a uniform mixture of active cells and passive dead ones—predicts the active density *in homeostasis* well and captures its relation to cell removal (degradation) and death rates (Fig. 4). The continuum model also provided a typical length scale beyond which cells are mechanically screened from the rest of the system. In a competition scenario, this screening length, combined with the fact that the system is never well-mixed at the cellular scale, leads to a specific growth advantage for the species with the higher active density near interfaces between the two species. This was here proposed to act through an ‘opportunistic growth’ mechanism that causes a quicker take-over of vacated space.

By having identical pressure dependencies for growth and identical death rates, the species in our system were originally designed such that the active density could be tuned without affecting homeostatic pressure in order to uncover the specific role of dead matter. However, fluctuations in our agent-based model did in fact lead to variations in homeostatic pressure across parameters (Fig. 4), presenting an unexpected opportunity to test its predictions as well, as it was previously suggested to be a competition driver for systems where homeostasis is regulated by pressure [4, 16]. The latter was shown to explain the suppression of tumour growth under mechanical pressure [4, 27, 44, 46] and can reliably predict fitness in systems where an equation of state exists between the pressure and the *active* density [16, 47]. Such a relation between active density and pressure implies that the predictions of both homeostatic properties—pressure and density—are equivalent. However, we saw that, when passive matter is continuously produced (and degraded) and growing cells are no longer the only mechanical component, these homeostatic properties are decoupled. The added passive matter of course does not rule out an equation of state, but if one exists it is between pressure and the *total* density rather then *active* density. Since competition outcomes were clearly correlated with increased active density, but not with homeostatic pressure (Fig. 5), our results do not only provide evidence for homeostatic density as an orthogonal competition driver, but also show that other competition drivers such as homeostatic pressure can be, at least to some extent, overpowered by it. Therefore, one could hypothesize that fast removal of dead/non-viable cells could be an important survival mechanism when competing against either rival species or mutations, motivating why, under certain circumstances, processes such as cell extrusion in epithelial tissues might be preferred to slow cell structure degradation. However, it should be noted that the balance between different competition drivers is parameter dependent. For example, in simulations with a significantly decreased friction coefficient *ζ*, we find that competition outcomes show a strong correlation with differences in homeostatic pressure, but not with those in active density (data not shown)—the opposite of the behavior seen in Fig. 5C,D). This is in line with an increased screening length *l_s_* for lower *ζ* and a resulting weaker competitive advantage through the opportunistic growth mechanism, as indicated by our theoretical considerations.

The simulations in this study were performed in 1D for simplicity and to distill the relevant mechanism by avoiding emergent collective phenomena arising e.g. from anisotropic cell shape, but can easily be generalized to higher dimensionality. Though highly idealized, real-world systems resembling this 1D single-file setup do in fact exist, such as bacteria growing in the “mother machine” [45, 48, 49], Eukaryotic cells in a capillary [50], and *Staphylococcus aureus* invasion of narrow bone canals (canaliculi) via fission [51]. In order to address the diversity of real biological systems, future work will have to examine the applicability of the uncovered mechanisms to higher dimensions, where interfaces between domains of different species do not have to be sharp and where higher dimensional collective effects come into play such as nematic order [52–54], cellular jamming [55] and intermittent load bearing force chains [56–58].

On a final note, we wish to point out that, while the rate at which forces change upon cell removal can potentially be arbitrarily fast in our model (for *E* → ±∞), the forces nevertheless vary continuously with time. To the best of our knowledge, this is the first numerical agent-based model of proliferation and removal in dense multicellular systems (even beyond the context of competition) where forces change continuously without any force jumps, throughout the life-cycle of the cell; i.e. both during cell death and removal and during division (see appendix B). Other numerical models (ranging from agent-based models [25, 47, 59] to active vertex models [26], cellular extended Potts models [60–63]) and the Subcellular Elements Method [64]), typically employ instantaneous cell division and/or death for computational simplicity leading to discontinuous forces. We could only find two examples where either division *or* death were implemented without force jumps [65, 66], but not both. Since the rate at which forces change can affect system properties, as was shown in this paper for the fitness and steady-state pressure, we suggest that instantaneous removal (or division) should be used with care. This is especially true when modelling realistic systems, where these processes are never truly instantaneous, but instead can have a wide range of rates as mentioned in the introduction.

## IV. ACKNOWLEDGEMENTS

We would like to thank Suropriya Saha, Tunrayo Adeleke-Larodo, Aboutaleb Amiri, Benoît Mahault, and Jérémy Vachier for discussions, feedback, and suggestions. We acknowledge support from the Max Planck School Matter to Life and the MaxSynBio Consortium, which are jointly funded by the Federal Ministry of Education and Research (BMBF) of Germany, and the Max Planck Society.

## Appendix A Methods

## Appendix B Numerical model details

Here we provide a more detailed account of the numerical model that was described concisely in the main text. A summary of all the parameters used in the model can be found in table I. The model is described in 1D as it was used in the simulations, but can be easily generalised to higher dimension.

**TABLE I.** Parameter values used in the simulation.

| Parameter | Value | Description |
| --- | --- | --- |
| $R$ | 0.5 | Node radius |
| $\Delta t$ | $5 \cdot 10^{-4}$ | Time step |
| $\langle \Gamma \rangle$ | 0.1 | Mean of normal distribution for base growth rates $\gamma_i$ |
| $\sigma_\Gamma$ | 0.05 | Std of base growth rates' normal distribution |
| $t_h^{\text{sim}}$ | 2000 | Simulation time for single species runs |
| $t_{\text{compete}}^{\text{sim}}$ | 5000 | Simulation time for competition runs |
| $H$ | 100 | Strength of Hertzian interaction |
| $\lambda$ | 3.5 | Threshold at which growth is considered stopped $\frac{1}{2}\text{erfc}(\lambda) = 3.7\text{e}^{-07}$ |
| $P^{\text{stall}}$ | 3.16 | Pressure at which $G(P_i)$ drops to $\frac{1}{2}\text{erfc}(\lambda)$<br>(corresponding to a node overlap of $\delta r_{i^\alpha j^\beta} = 0.1$) |
| $\zeta_0, \zeta^\parallel, \zeta^{\text{axis}}$ | 1 | Drag coefficients |
| $k^{\text{death}}$ | 0.01 | Cell death rate |
| $T^{\text{removal}}$ | 10 | Full duration of cell removal process (independent of $E$) |
| $L_x$ | 2560 | Channel length |

### Notations

Each cell is modelled as a dumbbell composed of two nodes of radius *R* that have a (time-dependent) preferred separation. The nodes’ positions are given by *r_i^α^_* where *i, j* are cell indices and the *α, β* indices differentiate between the two nodes of a cell. Without loss of generality we designate a cell’s left node as negative(−) and its right node as a positive(+). The separation between any two nodes is given by *r_i^α^j^β^_* ≡ *r_j^β^_* – *r_i^α^_*. For forces between any two nodes, we use the convention that the 1^st^ & 2^nd^ indices specify the node on which the force is applied, whereas 3^rd^ & 4^th^ indices specify the node causing the force:

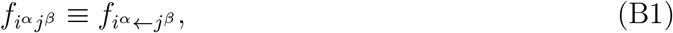

where the indices obey *i* ≠ *j* ∪ *α* ≠ *β*, since interactions can only happen between nodes (no self-propulsion of a node).

### Inter-cell interactions

The interaction between nodes belonging to different cells is a Hertzian repulsion:

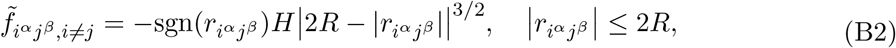

for any finite positive node overlap *δr_i^α^j^β^_* = 2*R* – *r_i^α^j^β^_*, and zero otherwise. Furthermore the following rule is applied:

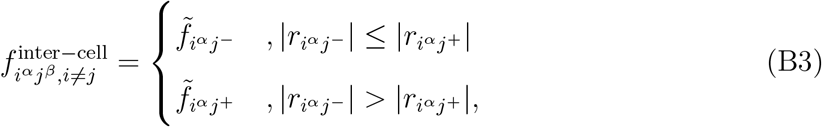

i.e. a given node *i^α^* interacts with *at most* one single node *j^β^* of any other cell *j* and specifically with the nearest of the two nodes of *j*. This choice makes physical sense, since overlaps of two cells are but a Hertzian approximation for deformations, and should not allow for interaction across physical objects (i.e. nodes). Yet another reason for this choice is force continuity as discussed in appendix C. When a cell dies and starts degrading, these forces are modulated as explained below in Eqs. (B6), (B8).

### Intra-cell forces, growth and division

As opposed to the inter-cell forces, the force between two nodes of the same cell can in principle be either repulsive or attractive (in order to keep structural integrity) and is given by a Hertzian “spring” with a (*growing*) rest-length 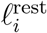:

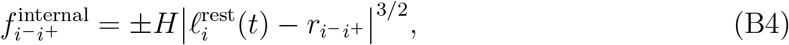

where *H* is a constant that provides the scale of the interaction strength, and the sign of the interaction is determined by 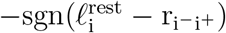. The similarity of this force-law to the inter-cell repulsion of Eq. (B2) is important for temporal force continuity as discussed in appendix C. The rest length 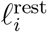 grows linearly with time from 0 (full overlap of the two nodes) immediately after division to a maximum of 2*R* (zero overlap). We express the growth of the cell equivalently via a growth phase 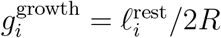 in the range [0, 1]. The growth rate of the cell is comprised of an inherent base growth rate *γ_i_* that is then modulated by pressure

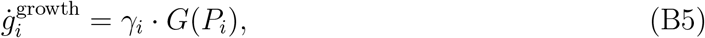

where in the 1D setup the pressure *P_i_* on a cell is simply the average of the external forces on it in the inward direction. The function *G*(*P_i_*) regulates the population growth in order to allow for stable homeostasis. The pressure dependence itself is identical for all cells, while the momentary value depends on the current pressure applied on the cell and obeys *G*(*P_i_* = 0) ≃ 1 and *G*(*P_i_* = *P*^sta11^) ≃ 0. Thus, for an isolated cell with no outside forces acting on it, the life-cycle time of a cell from division to division is 1/*γ_i_*. In this work we chose *G*(*P_i_*) to be a monotonously decreasing sigmoidal function as shown in Fig. 7 (the equation for *G*(*P_i_*) that appears in the main text is repeated in the figure caption for convenience). In all the simulations we use *λ* = 3.5 and a stall pressure of *P*^sta11^ ≃ 3.16, corresponding to the pressure felt by a cell if it is squeezed by its neighbors with an overlap of 10% the node diameter.

**FIG. 7.**
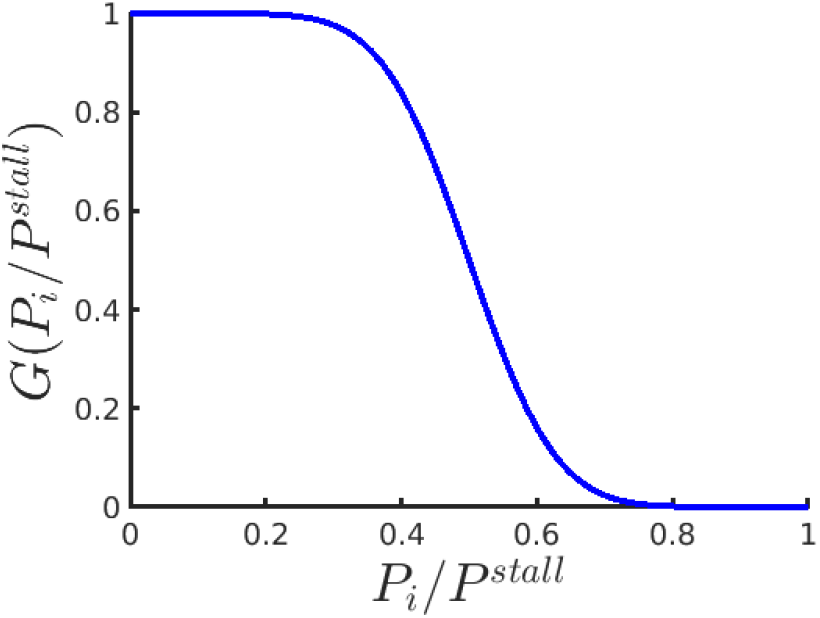
The pressure dependence 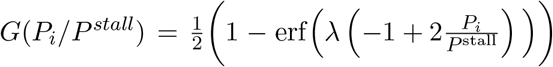 for the growth rate 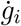 of cells, with *λ* = 3.5 and *P*^sta11^ ≃ 3.16.

### Division

If a cell *m* avoids death for long enough and its growth phase 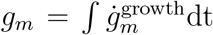 reaches 1 the cell divides, at which point each of the mother cell’s two nodes becomes a new cell, itself composed of two overlapping nodes. Each of the new cells *d*_1_, *d*_2_ starts with a growth phase 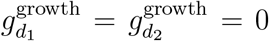. The base growth rates of the cells are drawn from a normal distribution Γ (compatible e.g. with bacterial phenotypic growth variability [32, 33, 67]) with a mean of 0.1 and a standard deviation 0.05. Additionally, when a partial mother-daughter inheritance scheme for the base growth rates was attempted (not shown), in line with e.g. bacterial proliferative self-regulation [33, 68, 69], only small quantitative differences are evident which do not change any of the conclusions presented in this paper.

### Degradation process of dying cells

Cell death is taken to be completely stochastic with a fixed rate *k*^death^ per cell. Once cell death is initiated, the cell irreversibly stops growing and cannot divide, transforming it from an To model a continuous degradation process each cell is assigned a scaling factor *S_i_* that modulates the forces it can exert on its neighbors as defined by Eqs. (B2), (B3), such that the net force on a given *node* is:

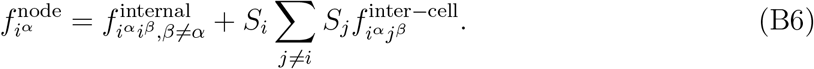

The force scaling factor is simply 1 as long as the cell is alive. Immediately after death a degradation phase 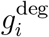 starts to accumulate according to which the scaling factors decay and finally vanish. The functional form of the scaling factor *S_i_* is here decaying exponentially:

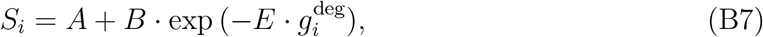

which together with the boundary conditions 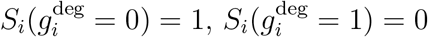 leads to:

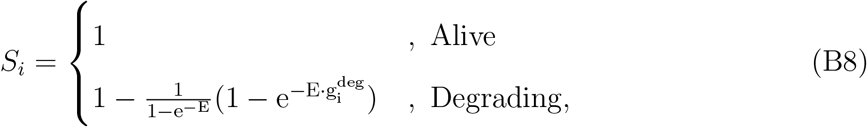

as shown in Fig. 8. The rate of cell degradation *k*^deg^ that is mentioned in the main text is taken as the inverse of the time from death initiation (when *S_i_* = 1) until the force scaling factor *S_i_* reaches a value of 0.1 (see Fig. 3)

**FIG. 8.**
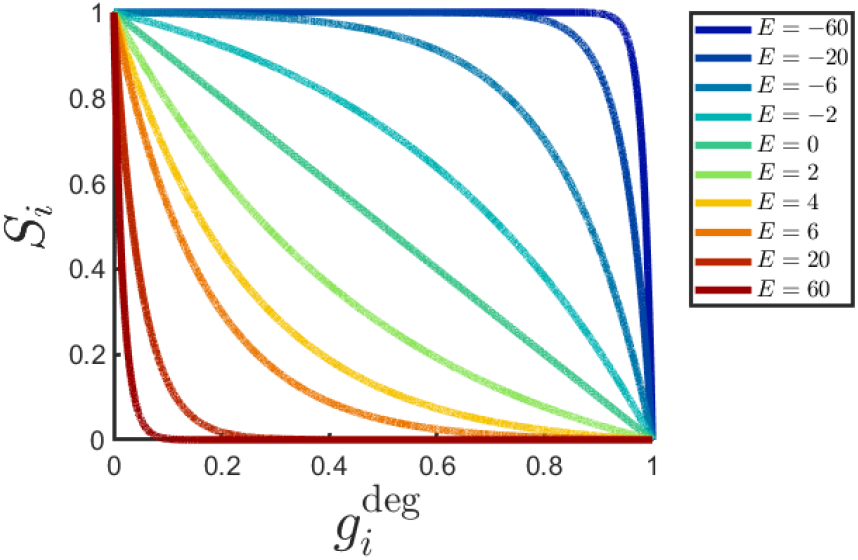
The functional form of the force scaling factors *S_i_* for different *E* values. The colors go from blue to red with increasing E, while brightness is correlated with longer collapse times and smaller pressure fluctuations (see Fig. 9).

### Equations of motion

The forces on a *cell* (as opposed to the individual nodes) can be decomposed into a component acting on the center of mass and a component 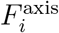 that drives the node separation degree of freedom:

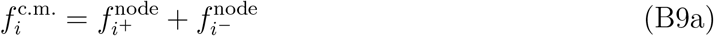

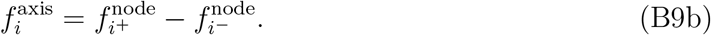

This leads to the following over-damped (zero Reynolds number limit) Newton equations for cell *i* in terms of its center of mass velocity 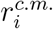 and the time derivative of the axial node-separation 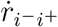:

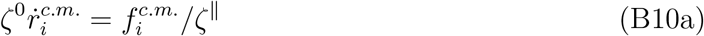

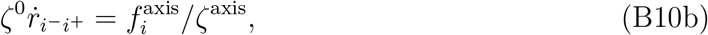

where *ζ*^0^ is the overall drag factor, while *ζ*^∥^ and *ζ*^axis^ define the relative contributions of the translational drag (parallel to cell long axis), and the axial stretch drag, respectively. Note that these equations do not contain self-propulsion nor any explicit noise terms. Thus single-cell diffusion is completely excluded and the only activity and stochasticity are in the growth, division and death which in turn drive the dynamics.

## Appendix C Continuity of forces during population changes

As mentioned in the discussion, the numerical model was constructed carefully in a way that avoids force discontinuities during cell proliferation, death and removal. Aside from the gradual cell degradation that is discussed extensively throughout the paper, two more mechanism are put in place to avoid force jumps during cell division. First, the rule in Eq. (B3) for the inter-cell forces prevents force jumps that would occur due to the instantaneous doubling of nodes upon cell division. Imagine a cell *m* is about to divide, while its node *α* has an overlap *δr_m^α^j^β^_* with the node of another cell *j*. When cell *m* divides, node *α* suddenly turns to two overlapping nodes of a new cell *d*_1_, both of which have the same overlap as before with the node of cell *j* and should thus apply double the force previously applied on that node of cell *j*. This is prevented by the rule in Eq. (B3).

The second measure for avoiding force jumps upon division, is replacing the growing rigid shape mechanism that is often used for modelling cell growth [59, 70], by a (non-linear) spring with a growing rest length and then matching the inter-cell force-law of Eq. (B2) to the intra-cell force law of Eq. (B4) that one gets just prior to division when 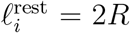. Thus the functional form of the node-node interactions doesn’t suddenly change when the nodes switch identity (from belonging to the same mother cell, to belonging to two distinct daughter cells) and neither the forces nor their derivatives jump.

These choices together with the explicit degradation process mean that all the forces on a cell change continuously, without any jumps, throughout the cell life-cycle. The exception is of course in the limits *E* → ±∞. In practice of course one also has to make sure the time step Δ*t* is small enough since *S_i_* can change quite sharply for very positive and very negative *E* values (see Fig. 8). Another benefit of the gradual removal mechanism is the added realism of the system staying confluent which can be obtained for low enough *E* values (Supplementary Video 1).

## Appendix D Typical length scale in homeostasis

In the continuum picture, growing active matter follows a continuity equation like (1), where we assume that the division and death rates *k*^div^ and *k*^death^ have suitable dependencies on the local mechanical environment (e.g., pressure or density), such that homeostasis can be reached. Without loss of generality, we assume that the net growth rate *α*(*P*) = *k*^div^(*P*) – *k*^death^(*P*) depends on *P*. For simplicity, we also consider a well-mixed system characterized by a single density *ρ*, such that

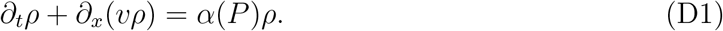

This equation has a spatially homogeneous steady-state *ρ* ≡ *ρ*^h^ defined by the condition *α*(*P*^h^) = 0 corresponding to homeostasis, where *P*^h^ = *P*(*ρ*^h^) via a constitute relation for the pressure. In real systems, usually *P* ≡ 0 below some critical density *ρ_c_* which corresponds to the closely packed (but uncompressed) density of cells and its functional form above *ρ_c_* is determined by the mechanical properties of the cellular material. While the specific form of *P*(*ρ*) is not important here, for consistency we have to require that *ρ*^h^ > *ρ_c_*, such that in its vicinity

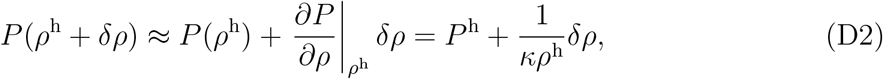

where *κ* is the compressibility of the cellular material at homeostatic density.

To achieve stability, we also require that

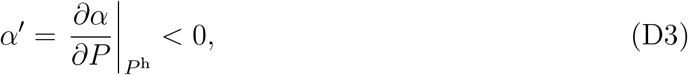

as this ensures that, again near *ρ*^h^, the net growth rate

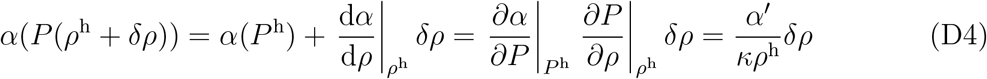

opposes *δρ*. Local force balance in this over-damped system reads

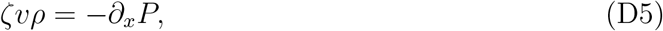

where *ζ* is the friction coefficient with the environment or substrate. Plugging relations (D2), (D4) and (D5) into equation (D1), we arrive at the following equation for a perturbation *ρ* = *ρ*^h^ + *δρ* of the homogeneous, homeostatic state:

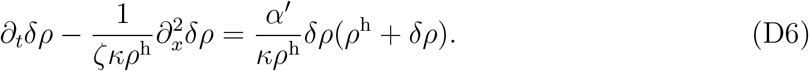

Neglecting higher order terms, we have

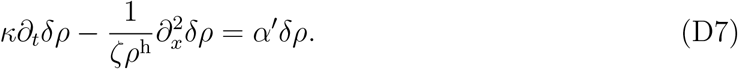

Perturbations therefore both spread diffusively by mechanical relaxation with a diffusion constant *D* = 1/(*κζρ*^h^) and decay actively (through growth/degradation of cellular matter) with a rate *γ* = *α′/κ*. This defines a typical length scale 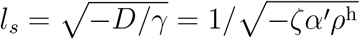 which perturbations are able to affect. Note that the independence of *l_s_* from *κ* is a consequence of *α* being defined as a function of *P* (and not, for example, *ρ* directly), such that both mechanical relaxation and growth adaptation scale equally with *κ*. For the value of the drag coefficient in our simulations (see Table I), a total homeostatic density of order 0.8 (see, e.g., the right end of Fig. 4A) and *α′* ≈ ln(2)〈Γ〉*G′*(*P*^h^) ≈ −0.044 with 〈Γ〉 = 0.1 (Table I) and *P*^h^ ≈ 1.95 (Fig. 4B), this length scale evaluates to *l_s_* ≈ 5.3 in our simulations.

*l_s_* can be interpreted as a screening length: Consider a continuous external perturbation (for example, a density sink) which fixes *δρ*(*x* = 0) = *δρ*_0_ at the boundary of 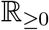. Then, the steady state of equation (D7) is given by *δρ*(*x*) = *δρ*_0_ exp(−*x/l_s_*), indicating exponential screening of the external perturbation. An increased friction *ζ* with the substrate (lower mobility) or a stronger change in the growth-degradation balance for a local deviation from the homeostatic pressure (*α′*) will therefore lead to a stronger attenuation away from the perturbation and consequently a shorter *l_s_*. Conversely, in the limit of large mobilities (small friction coefficient *ζ*) or slow growth responses *α′*, the screening length diverges and the system approaches the mean-field description in which pressure/density equilibration is the fastest process.

## Appendix E Interface dynamics in a simple lattice model

We would like to examine the possible influence of the finite granularity of the system on competition dynamics. By granularity we mean the fact that the system is comprised of discrete units, which can be either passive (dead) or active (have the potential to divide). In the main text, we showed that an entirely mean-field, continuum consideration with well-mixed active and passive matter predicts no particular advantage for a cell species which produces more or less passive matter. However, we also showed that, in reality, there exists a finite screening length in the system, so that growth in response to a local perturbation in pressure happens only in a finite region around a perturbation (see appendix D). If this length scale is on the order of the cell size, consequently, only a finite number of discrete cells will respond to a perturbation (for example, the removal of a cell), possibly triggering small-number effects. To investigate the possible consequences of this granularity, we consider the limit of very small screening lengths, in which only neighbors are able to contribute significantly to growth in response to a pressure drop.

To this end, we construct a stochastic lattice model which is characterized by only two different rates: *k*^death^ and *k*^deg^, the rate at which an active cell turns into a passive (non-dividing) cell, and the rate at which passive cells are removed from the system, respectively. The role of the growth-limiting mechanical environment (e.g., through pressure) in this model is played by the finite availability of lattice sites, where growth can only occur if a lattice site is freed up by cell removal. This also conserves the property of both the particle-based and the continuum model, where the homeostatic density/pressure do not depend on composition, as the growth behavior of all active cells is identical, irrespective of whether they belong to a species that leaves more or less dead matter behind.

In a single species, lattice sites will be occupied by a mixture of active cells that become passive with rate *k*^death^ and passive cells that are removed with rate *k*^deg^. Once a cell is removed, one of the nearby active cells immediately divides and fills the gap (we will clarify below what exactly we mean by “nearby”). In steady-state, when the total number of cells is constant, the fraction of active cells is therefore

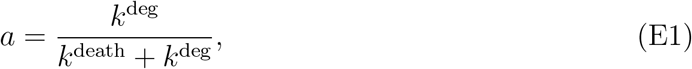

which is the discrete equivalent of Eq. (4). At the same time, this is the fraction of time the average cell spends in its active state, or the probability of finding a randomly picked cell to be in an active state.

We consider a one-dimensional configuration where each half-space is filled by a different species. Similar to the particle-based model that we consider later in the main text, they only differ in the rate *k*^deg^ with which passive cells are removed. Without loss of generality, we assume that the species on the left has a high degradation rate 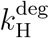 compared to the species on the right with 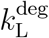, which implies a higher equilibrium active fraction *a*_H_ > *a*_L_ as well.

What then happens at the interface, where a cell of the left species directly neighbors a cell from the right species? First, we note that in the event of the removal of one of the two central cells, a symmetrical distribution of species arises: The gap is lined by H cells to its left and L cells to its right. Since we consider two species with identical growth dynamics (for the active cells), we also assume that it is only the number and position of the active cells of each species that determine which species will fill the gap. Let 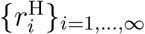 be the distances of the *active* cells of species H, numbered in increasing distance from the interface, and 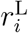 the corresponding distances for species *L*. Consistent with our screening length consideration, we assume a weighting function *w*(*r*) that determines the growth contribution of an active cell at distance *r* from the interface, with *w*(*r*) → 0 as *r* → ∞. Given a specific configuration characterized by 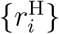 and 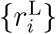, the probability that species H will fill the gap is then equal to its total relative growth contribution, i.e. 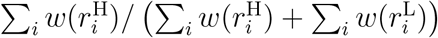. With this probability for a single given configuration, the probability of species H filling the gap in a large ensemble of such events with arbitrary configurations is

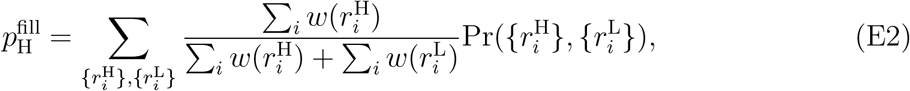

where the sum goes over all possible configurations that have a non-zero total contribution and 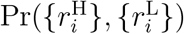 is the probability of the configuration 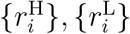 among these. For simplicity, we now assume a sharp cutoff where a maximum of *d* cells on either side of the interface can contribute to filling the gap (essentially a discrete equivalent of the screening length), i.e.

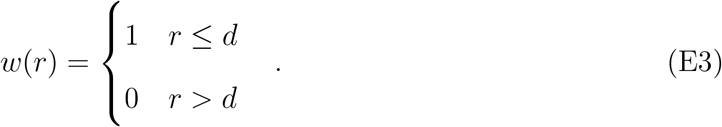

In this case, the sums over the weights in equation (E2) simply count the active cells in each half-space within distance *d* from the interface, and the probability of having a specific number of active cells, e.g. *k* on the left and *l* on the right, is binomially distributed, such that

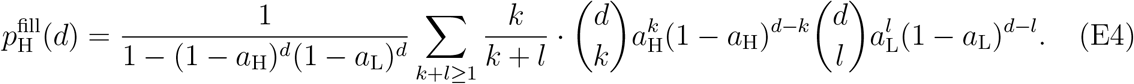

To estimate the maximum effect, we explicitly calculate 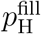 for *d* =1 from equation (E4), such that only the nearest neighbors determine growth:

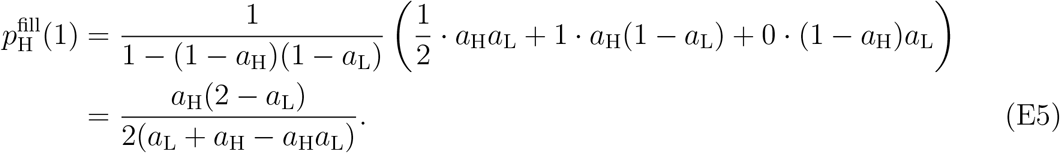

For large *d*, the relative numbers of active cells in each half space become increasingly more narrowly distributed around their expectation values *a*_H_*d* and *a*_L_*d*, so we can approximate Eq. (E4) as

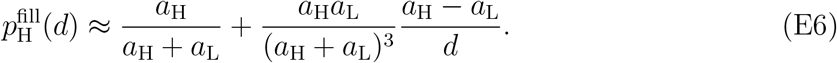

The second term corresponds to a finite-size correction with respect to the limit 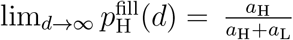, where exactly a fraction of *a*_H_ cells in the left half-space and a fraction of *a*_L_ cells in the right half space are active. This shows that, in any symmetrical finite sample of cells around an interface, the growth contribution from active cells is statistically biased in favor of the species with the higher active fraction *a*_H_. Note that the limit for infinite *d* should hold for any reasonable weighting function *w*(*r*) where *d* represents the scale on which cells contribute significantly to growth (not just Eq. (E3)).

Having determined the probability with which each side will fill the gap, we need to calculate the frequency with which gaps are created in the first place. Overall, species H will only have an advantage if, on average, gaps created by the removal of L cells are filled by H cells with a higher rate than gaps created by the removal of H cells are filled by L cells. Given a cell that is passive (which is the case with probability 1 – *a*), it is removed with rate *k*^deg^, leading to a total gap creation rate for a single site of

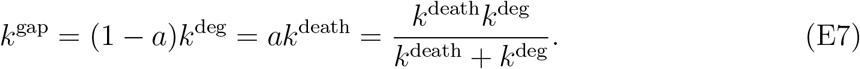

Consistently, for very large *k*^deg^ (i.e. no passive cells), we simply have *k*^gap^ = *k*^death^, whereas for small *k*^deg^, removal of passive cells is the bottleneck. Note that 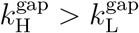, i.e. gaps are created more frequently on the side with a higher percentage of active cells. However, we have to consider both gap creation and cell replacement to calculate the net bias. Consequently, the total rate *k*_L→H_ with which cells of type L are replaced with cells of type H directly at the interface is

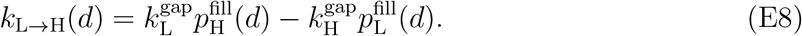

For the *d* = 1 case, this evaluates to

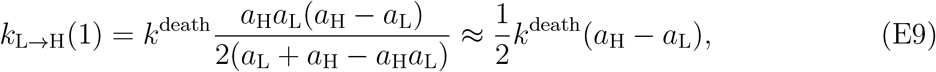

where the last step is valid for *a*_H_, *a*_L_ close to 1. Therefore, for *d* =1 (and, in fact, for any finite *d*), the lower 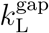 with which species L opens a gap is more than outweighed by species H’s statistically increased probability to fill it. This creates a bias towards species H which has a higher active fraction *a*_H_. Note that by using equation (E6) we can see that for large *d*,

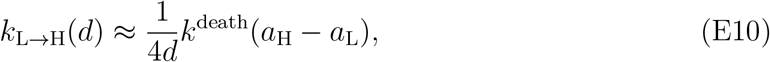

and consequently,

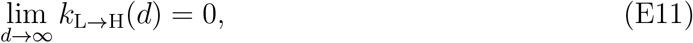

which underscores the nature of this bias as a small-number effect due to the granularity of the system in conjunction with the screening length.

It should be noted that the correspondence between the true, agent-based model (or a continuum model) and this lattice model should be regarded as conceptual rather than formal. While *d* above does play the role of a screening length in the sense that it allows only cells within a certain distance of the interface to replenish the system after the removal of an interface cell, there are many differences which forbid the interpretation of these lattice model results as a quantitative prediction. Most important among these are the instantaneous (rather than continuous) growth and the absence of explicit mechanical relaxation (which is only incorporated in an effective sense through the screening length). For example, with a more continuous growth process in mind, the probability *p*_H_ of filling the gap with a cell of type H should rather be viewed as the fraction of the gap which is filled with cellular material of type *H*. Therefore, this analysis should be interpreted as “supporting evidence” for our hypothesis that a finite screening length in conjunction with the discreteness of cells can lead to a biased selection of active cells from the more active species, rather than a formal proof.

## Appendix F Local and global pressure fluctuations

As already pointed out in the main text, the homeostatic pressure measured in the simulations (Fig. 4B) shows a deviation from the constant value predicted by the zeroth-order mean-field approximation 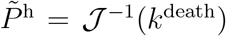, where 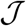 defines the pressure dependency of the division rate, 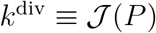. The reason for this is the degradation of cells, which entails a constant turnover of matter. Due to the screening length, degrading cells cause *local* deviations from the mean-field uniform pressure derived in the continuum description in the main text.

First, let us derive this zeroth-order prediction of the homeostatic pressure (i.e. where division and death exactly balance) in the context of the agent-based numerical model: In homeostasis, the probabilities for death and for survival until division have to be equal. Cells die with a constant rate *k*^death^, leading to an exponentially distributed time of death, such that the condition for the survival probability becomes *p*^survival^ = exp (−*k*^death^*τ*^div^) = 1/2, where *τ*^div^ is the resulting (pressure modulated) division time in homeostasis. This leads to 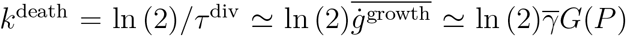, where 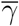 is the average base growth rate (which we approximate below with the the mean of the distribution of growth rates for new cells Γ). Inverting *G* one gets for the homeostatic pressure

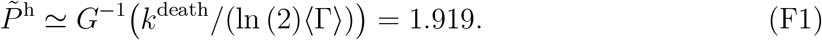

Second, we examine how the time-averaged balance between the division rate and the death rate is modified by local pressure fluctuations *P* = *P*^h^ + *δP* about the unknown homeostatic pressure *P*^h^, given the nonlinear dependence of division on pressure through 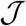. For this purpose, we expand the time-averaged division rate

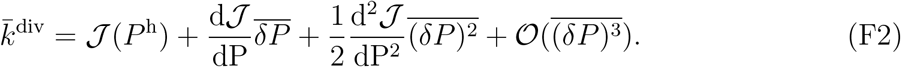

The term linear in *δP* drops out in the time averaging and extracting the homeostatic pressure we get

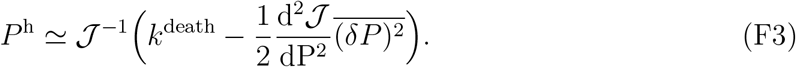

Since 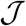 is a decreasing function of pressure and 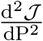 is positive (see Fig. 7 with 0.61 ≤ *P*^h^/*P^stall^* ≤ 0.62 in simulation), we see that, for larger local fluctuations, the pressure increases (if one can neglect the the change of 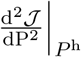 itself).

With this expectation in mind, it is worth re-examining Fig. 4B. If local pressure fluctuations are small, we expect a homeostatic pressure equal to the zeroth-order prediction (F1) indicated by the dashed orange line. The fact that the homeostatic pressure is always above this value (the minimum value measured being 1.924) indicates that the fluctuations are always relevant for the chosen cell removal mechanism and parameters used. Furthermore, we expect that the fluctuations would be smallest at around *E* = 4 where the measured homeostatic pressure is minimal, and for the fluctuations to increase away from this value towards both higher and lower *E* values [71].

Indeed, looking at the *global* pressure fluctuations in Fig. 9A, we see that they are minimal at around the same *E* value. Note that a *global* pressure fluctuation represents, at any point in time, the sum of contributions of individual *local* pressure fluctuations with only weak spatial correlations beyond the screening length. Thus, *global* fluctuations should decay 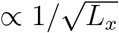 for system sizes *L_x_* much larger than the screening length. This is indeed the case as shown in Fig. 10B.

**FIG. 9.**
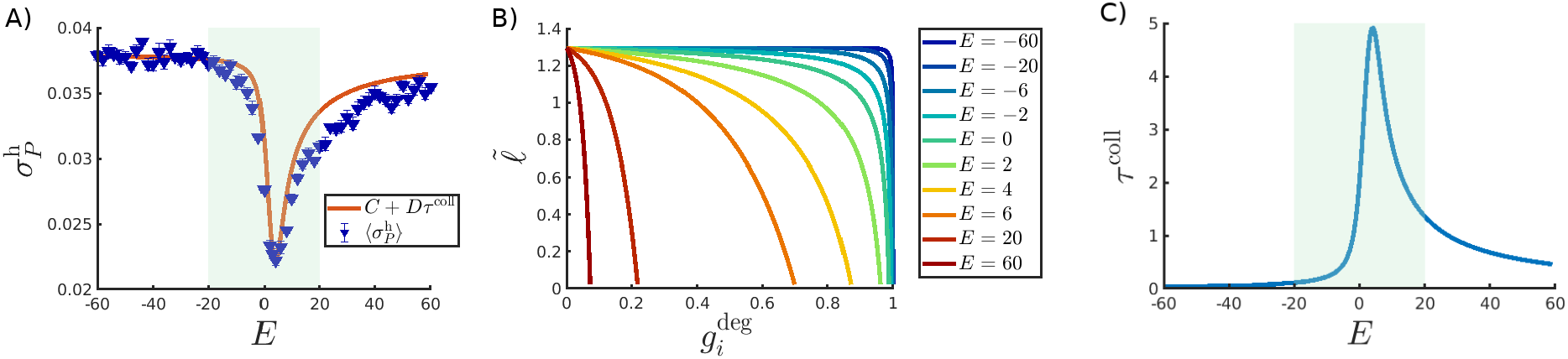
A) Global pressure fluctuations. Blue triangles denote the standard deviation of the pressure measurement in a given simulation (averaged over 9-10 realizations). The orange line depicts a linear transformation of the collapse time scale shown in panel C (using the coefficients *C* = 0.38, *D* = −0.0032). B) The mean-field approximation for the cell length as a function of the degradation phase. Same color scheme as in Fig. 8. For this figure we used the minimal pressure value measured in the simulation *P*^ref^ = 1.965 and 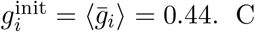) The time scale of cell mechanical ‘‘collapse” defined in Eq. (F5) (using the same 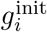 as in panel B).

**FIG. 10.**
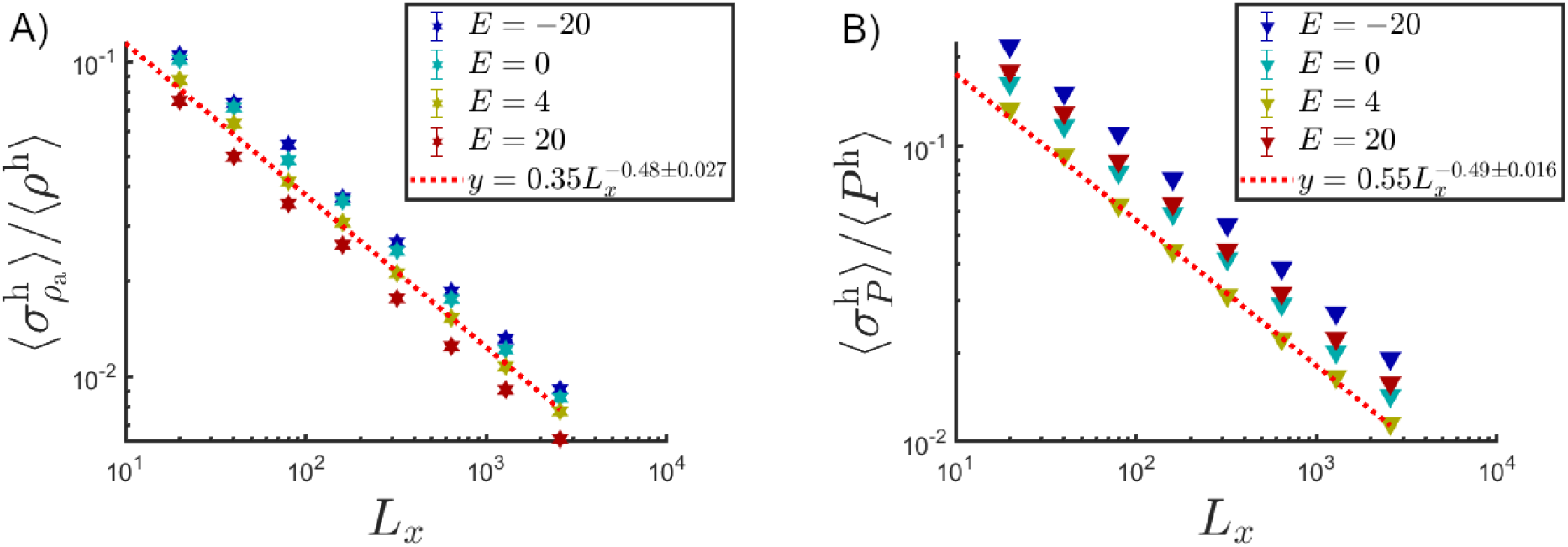
Standard deviation of the fluctuations in the homeostatic active density (A) and in the homeostatic pressure (B) normalized by the mean rate as a function of system sizes *L_x_* for various values of the softening factor *E*. Each data point is averaged over 10 realizations (error bars are smaller than the size of the markers). The red dashed lines are power-law fits to the *E* = 4 data set. The exponent is in both cases −0.5 to within the measurement error, as expected for localized uncorrelated fluctuations and meaning that finite-size effects can be neglected.

Now that we know that, up to said 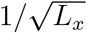 scaling, *global* pressure fluctuations are a good proxy for fluctuations of the average pressure on length scales of the screening length, we would like to show how changes in E, i.e. the dynamics of the degradation mechanism, affect the latter *local* pressure fluctuations. For this purpose, we consider the change in length of a single degrading cell with time. As an approximation, we start with the rest length of a cell and modulate it by the decaying force scaling factor *S_i_* while keeping the effects of the surrounding cell medium fixed (such as the pressure applied by it on the cell, *P*^ref^):

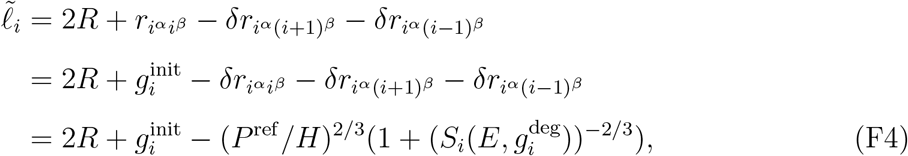

where for the third equality the overlaps are calculated using Eqs. (B2)-(B4), (B6) which give 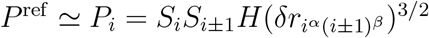 and 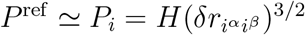. Here 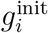 is the growth phase at which the cell died and started degrading. Examples of the decay of this length with time are shown in Fig. 9B for several *E* values (using representative values for 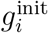, *P*^ref^). For very negative *E* values, the cell length decays only slowly for most of the degradation time before finally dropping sharply. We call this sharp drop in length the ‘collapse’ stage and associate to it a large local pressure drop. Such collapse could describe for example the loss of structural integrity due to tearing of the cell membrane in eukaryotes or breaking of the cell wall in bacterial lysis [24]. For very high *positive E* values, the cell starts collapsing as soon as it dies. As a measure of the collapse time scale *τ*^coll^ we consider the time it takes the length of a degrading cell to decay from 0.9 to 0.1 of its initial value (when *S_i_* = 1):

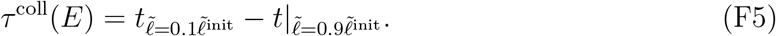

Of course, this measure for the collapse time scale depends on the growth phase of a cell when it died 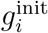, but we verified that, for our model, it only does so weakly. The collapse time scale is shown in Fig. 9C. It is reasonable to associate faster cell collapse (small *τ*^coll^) with larger *local* pressure fluctuations. Indeed, a simple linear transformation of *τ*^coll^ matches the measured *global* pressure fluctuations as shown in Fig. 9A, which we showed above to be a proxy for the *local* fluctuations. This suggests that pressure fluctuations in our system are indeed determined by the characteristics of the degradation process.

## Appendix G Competition dynamics at short and long time scales

We remind the reader that the original simulation time of 5000 for the competition assay was chosen to be short enough to avoid measurement bias caused by the absorbing boundary (where an entire sample is taken over by a single species). In Figs. 11A,11B we confirm that the competition does not saturate at some finite species fraction, but that indeed the less fit species is eradicated from the system at the long-time limit.

**FIG. 11.**
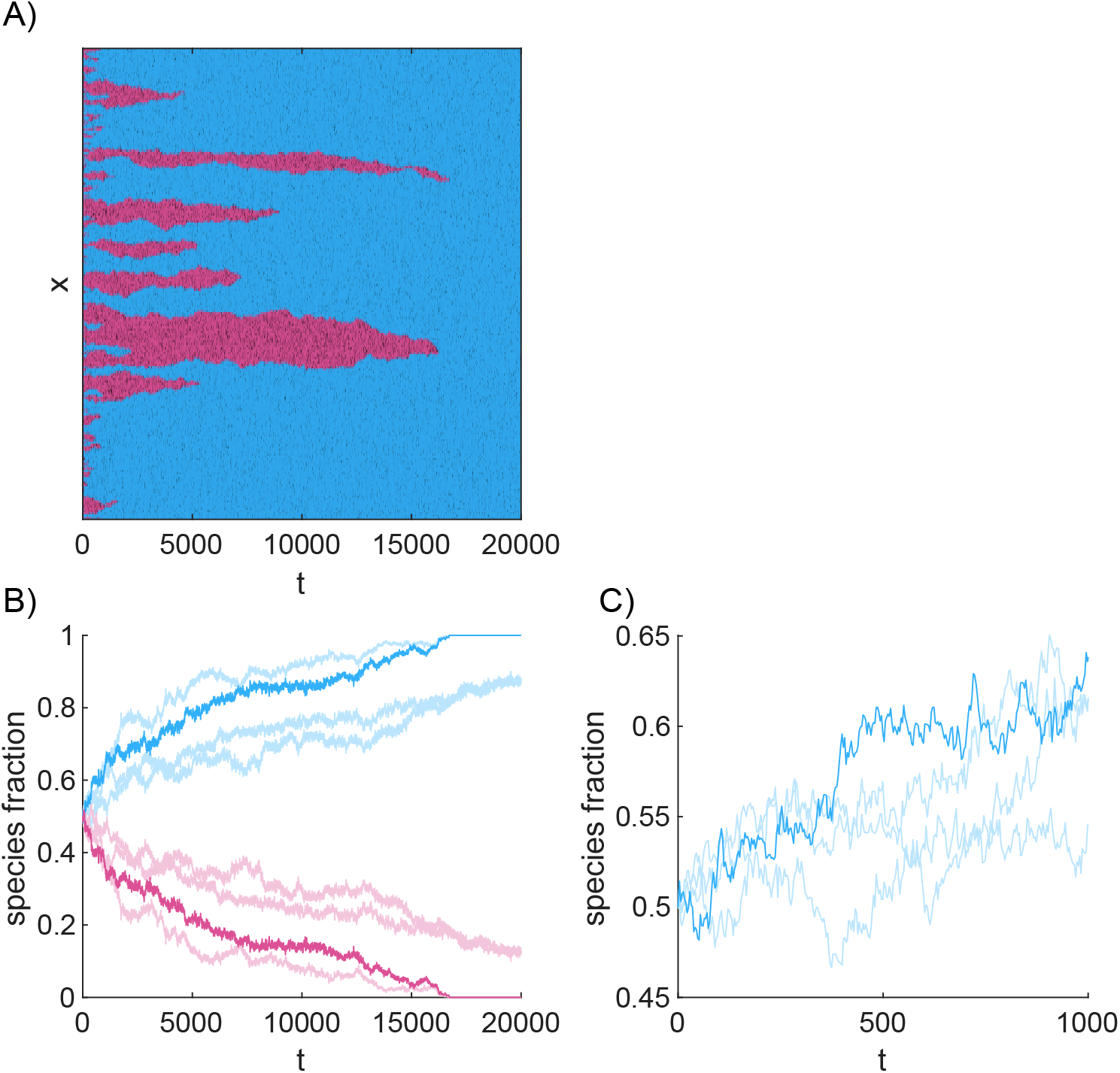
A) Kymograph showing the space-time evolution of two competing species with a large difference in homeostatic active density (*E*_H_ = 16, 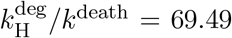 *E*_L_ = −20, 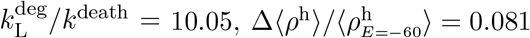) in simulations that were 4 times as long as the ones in the main text (*t*^sim^ = 20,000). System size is 4 times smaller than the one used in the main text (*L_x_* = 640) in order to be able to show the entire system in one figure without obscuring too much details in the visual coarse-graining. On a long enough time scale one species fully takes over the available volume and the other becomes extinct. B) Time series of the species fraction for 4 realizations of the same setup and parameters. The bolder line corresponds to the specific realization depicted in panel A. On this time scale 2 out of 4 realizations shown extinction of species L. C) Zoom in on the early evolution of the simulations used in panel B (only showing species H). The stochastic fluctuations can cause the H species to temporarily have a lower fraction than species L (fraction < 0.5), and still out-compete species L in the long-term.

It is also useful to look at the species fraction dynamics at start of the competition assay as shown in Fig. 11C. One can see that the fraction of species H can go below 0.5 due to the stochastic nature of the dynamics, giving way to species L, before increasing again and eventually winning. This shows that the dynamics is robust to the choice of initial conditions and that an initial larger fraction of L cells will not change the identity of the winner.

## References

[1] D. J. Jörg, Y. Kitadate, S. Yoshida, and B. D. Simons, Stem cell populations as self-renewing many-particle systems, Annual Review of Condensed Matter Physics 12 (2020).

[2] B. I. Shraiman, Mechanical feedback as a possible regulator of tissue growth, Proceedings of the National Academy of Sciences 102, 3318 (2005).

[3] S. Moitrier, C. Blanch-Mercader, S. Garcia, K. Sliogeryte, T. Martin, J. Camonis, P. Marcq, P. Silberzan, and I. Bonnet, Collective stresses drive competition between monolayers of normal and ras-transformed cells, Soft matter 15, 537 (2019).

[4] M. Basan, T. Risler, J.-F. Joanny, X. Sastre-Garau, and J. Prost, Homeostatic competition drives tumor growth and metastasis nucleation, HFSP journal 3, 265 (2009).

[5] M. E. Hibbing, C. Fuqua, M. R. Parsek, and S. B. Peterson, Bacterial competition: surviving and thriving in the microbial jungle, Nature reviews microbiology 8, 15 (2010).

[6] J. Friedman, L. M. Higgins, and J. Gore, Community structure follows simple assembly rules in microbial microcosms, Nature Ecology & Evolution 1 (2017).

[7] B. Drossel and M. Kardar, Phase ordering and roughening on growing films, Phys. Rev. Lett. 85, 614 (2000).

[8] R. Levayer and E. Moreno, Mechanisms of cell competition: themes and variations, Journal of cell biology 200, 689 (2013).

[9] C. Clavería and M. Torres, Cell competition: mechanisms and physiological roles, Annual review of cell and developmental biology 32, 411 (2016).

[10] L. Speare, A. G. Cecere, K. R. Guckes, S. Smith, M. S. Wollenberg, M. J. Mandel, T. Miyashiro, and A. N. Septer, Bacterial symbionts use a type vi secretion system to eliminate competitors in their natural host, Proceedings of the National Academy of Sciences 115, E8528 (2018).

[11] C. Darwin, On the origin of species (London: J. Murray, 1859).

[12] G. F. Gause, The struggle for existence (Baltimore: Williams & Wilkins, 1934).

[13] L. Wagstaff, M. Goschorska, K. Kozyrska, G. Duclos, I. Kucinski, A. Chessel, L. Hampton-O’Neil, C. R. Bradshaw, G. E. Allen, E. L. Rawlins, et al., Mechanical cell competition kills cells via induction of lethal p53 levels, Nature communications 7, 1 (2016).

[14] M. Danino, D. A. Kessler, and N. M. Shnerb, Stability of two-species communities: drift, environmental stochasticity, storage effect and selection, Theoretical population biology 119, 57 (2018).

[15] S. Chu, M. Kardar, D. R. Nelson, and D. A. Beller, Evolution in range expansions with competition at rough boundaries, Journal of theoretical biology 478, 153 (2019).

[16] M. Basan, J. Prost, J.-F. Joanny, and J. Elgeti, Dissipative particle dynamics simulations for biological tissues: rheology and competition, Physical Biology 8, 026014 (2011).

[17] E. Marinari, A. Mehonic, S. Curran, J. Gale, T. Duke, and B. Baum, Live-cell delamination counterbalances epithelial growth to limit tissue overcrowding, Nature 484, 542 (2012).

[18] G. T. Eisenhoffer, P. D. Loftus, M. Yoshigi, H. Otsuna, C.-B. Chien, P. A. Morcos, and J. Rosenblatt, Crowding induces live cell extrusion to maintain homeostatic cell numbers in epithelia, Nature 484, 546 (2012).

[19] A. Gelimson and R. Golestanian, Collective dynamics of dividing chemotactic cells, Phys. Rev. Lett. 114, 028101 (2015).

[20] A. Matamoro-Vidal and R. Levayer, Multiple influences of mechanical forces on cell competition, Current Biology 29, R762 (2019).

[21] T. B. Saw, A. Doostmohammadi, V. Nier, L. Kocgozlu, S. Thampi, Y. Toyama, P. Marcq, C. T. Lim, J. M. Yeomans, and B. Ladoux, Topological defects in epithelia govern cell death and extrusion, Nature 544, 212 (2017).

[22] B. Loewe, M. Chiang, D. Marenduzzo, and M. C. Marchetti, Solid-liquid transition of deformable and overlapping active particles, Physical Review Letters 125, 038003 (2020).

[23] Y. Gavrieli, Y. Sherman, and S. A. Ben-Sasson, Identification of programmed cell death in situ via specific labeling of nuclear dna fragmentation., Journal of cell Biology 119, 493 (1992).

[24] L. Turnbull, M. Toyofuku, A. L. Hynen, M. Kurosawa, G. Pessi, E. S. Gloag, R. Shimoni, U. Omasits, N. K. Petty, S. R. Osvath, G. Ca, S. Ito, X. Yap, L. G. Monahan, R. Cavaliere, C. H. Ahrens, I. G. Charles, N. Nomura, L. Eberl, and C. B. Whitchurch, Explosive cell lysis as a mechanism for the biogenesis of bacterial membrane vesicles and biofilms, Nature Communications (2016).

[25] F. Jülicher, T. Bittig, O. Wartlick, A. Kicheva, and M. González-Gaitán, Dynamics of anisotropic tissue growth, New Journal of Physics (2008).

[26] D. L. Barton, S. Henkes, C. J. Weijer, and R. Sknepnek, Active Vertex Model for cell-resolution description of epithelial tissue mechanics, PLOS Computational Biology 13, 1 (2017).

[27] F. Montel, M. Delarue, J. Elgeti, D. Vignjevic, G. Cappello, and J. Prost, Isotropic stress reduces cell proliferation in tumor spheroids, New Journal of Physics 14 (2012).

[28] N. Minc, A. Boudaoud, and F. Chang, Mechanical forces of fission yeast growth, Current Biology 19, 1096 (2009).

[29] A. Puliafito, L. Hufnagel, P. Neveu, S. Streichan, A. Sigal, D. K. Fygenson, and B. I. Shraiman, Collective and single cell behavior in epithelial contact inhibition, Proceedings of the National Academy of Sciences 109, 739 (2012).

[30] J. Ranft, M. Aliee, J. Prost, F. Jülicher, and J.-F. Joanny, Mechanically driven interface propagation in biological tissues, New Journal of Physics 16, 035002 (2014).

[31] D. Drasdo, S. Hoehme, and M. Block, On the role of physics in the growth and pattern formation of multi-cellular systems: what can we learn from individual-cell based models?, Journal of Statistical Physics 128, 287 (2007).

[32] A. Amir, Cell size regulation in bacteria, Phys. Rev. Lett. 112, 1 (2014).

[33] J. Lin and A. Amir, The Effects of Stochasticity at the Single-Cell Level and Cell Size Control on the Population Growth, Cell Systems 5, 358 (2017).

[34] J. H. Irving and J. G. Kirkwood, The statistical mechanical theory of transport processes. iv. the equations of hydrodynamics, The Journal of Chemical Physics 18, 817 (1950).

[35] We checked that the use of different measures for calculating the homeostatic pressure, such as the pressure exerted on a wall or cell-sensed pressure, does not introduce any significant changes in the pressure measurement except possibly a fixed linear scaling and thus does not change any of the conclusions in this paper.

[36] Since a measure for the density of passive matter was not uniquely defined for the agent-based numerical model, one can instead take the mean-field limit discussed under “theoretical considerations” and use Eq. (4) to calculate the total density e.g. using 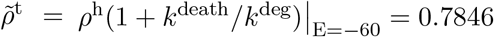.

[37] D. A. Kessler and N. M. Shnerb, The effect of spatial heterogeneity on the extinction transition in stochastic population dynamics, New Journal of Physics 11, 043017 (2009).

[38] S. Moitrier, Competition between normal and transformed cell populations., Ph.D. thesis, Universitáe Paris sciences et lettres (2017).

[39] R. J. Murphy, P. R. Buenzli, R. E. Baker, and M. J. Simpson, Mechanical cell competition in heterogeneous epithelial tissues, Bulletin of Mathematical Biology 82, 1 (2020).

[40] A. Bove, D. Gradeci, Y. Fujita, S. Banerjee, G. Charras, and A. R. Lowe, Local cellular neighborhood controls proliferation in cell competition, Molecular biology of the cell 28, 3215 (2017).

[41] D. Gradeci, A. Bove, A. R. Lowe, S. Banerjee, and G. Charras, Distinct modes of cell competition are governed by entropic and energetic properties of mixed cell populations, bioRxiv, 729731 (2019).

[42] C. Brás-Pereira and E. Moreno, Mechanical cell competition, Current opinion in cell biology 51, 15 (2018).

[43] M. J. Paszek, N. Zahir, K. R. Johnson, J. N. Lakins, G. I. Rozenberg, A. Gefen, C. A. Reinhart-King, S. S. Margulies, M. Dembo, D. Boettiger, et al., Tensional homeostasis and the malignant phenotype, Cancer cell 8, 241 (2005).

[44] R. Kaukonen, A. Mai, M. Georgiadou, M. Saari, N. De Franceschi, T. Betz, H. Sihto, S. Ventelä, L. Elo, E. Jokitalo, J. Westermarck, P. L. Kellokumpu-Lehtinen, H. Joensuu, R. Grenman, and J. Ivaska, Normal stroma suppresses cancer cell proliferation via mechanosensitive regulation of JMJD1a-mediated transcription, Nature Communications 7(2016).

[45] D. Yang, A. D. Jennings, E. Borrego, S. T. Retterer, and J. Männik, Analysis of factors limiting bacterial growth in pdms mother machine devices, Frontiers in Microbiology 9, 871 (2018).

[46] M. Delarue, F. Montel, D. Vignjevic, J. Prost, J. F. Joanny, and G. Cappello, Compressive stress inhibits proliferation in tumor spheroids through a volume limitation, Biophysical Journal 107, 1821 (2014).

[47] J. Ranft, M. Basan, J. Elgeti, J.-F. Joanny, J. Prost, and F. Julicher, Fluidization of tissues by cell division and apoptosis, Proceedings of the National Academy of Sciences 107, 20863 (2010).

[48] P. Wang, L. Robert, J. Pelletier, W. L. Dang, F. Taddei, A. Wright, and S. Jun, Robust growth of escherichia coli, Current Biology 20, 1099 (2010).

[49] Ö. Baltekin, A. Boucharin, E. Tano, D. I. Andersson, and J. Elf, Antibiotic susceptibility testing in less than 30 min using direct single-cell imaging, Proceedings of the National Academy of Sciences 114, 9170 (2017).

[50] S. H. Au, B. D. Storey, J. C. Moore, Q. Tang, Y.-L. Chen, S. Javaid, A. F. Sarioglu, R. Sullivan, M. W. Madden, R. O’Keefe, D. A. Haber, S. Maheswaran, D. M. Langenau, S. L. Stott, and M. Toner, Clusters of circulating tumor cells traverse capillary-sized vessels, Proceedings of the National Academy of Sciences (2016).

[51] K. L. de Mesy Bentley, R. Trombetta, K. Nishitani, S. N. Bello-Irizarry, M. Ninomiya, L. Zhang, H. L. Chung, J. L. McGrath, J. L. Daiss, H. A. Awad, S. L. Kates, and E. M. Schwarz, Evidence of staphylococcus aureus deformation, proliferation, and migration in canaliculi of live cortical bone in murine models of osteomyelitis, Journal of Bone and Mineral Research 32, 985 (2017).

[52] D. Dell’Arciprete, M. Blow, A. Brown, F. Farrell, J. S. Lintuvuori, A. McVey, D. Marenduzzo, and W. C. Poon, A growing bacterial colony in two dimensions as an active nematic, Nature communications 9, 1 (2018).

[53] Q. Zhang, J. Li, J. Nijjer, H. Lu, M. Kothari, R. Alert, T. Cohen, and J. Yan, Mechanical stress determines morphogenesis and cell ordering in confined bacterial biofilms, bioRxiv (2021).

[54] L. Hupe, P. Bittihn, Y. G. Pollack, A. Amiri, and R. Golestanian, Rods vs. dumbbells in growing active nematics, In preparation (in preparation).

[55] S. Garcia, E. Hannezo, J. Elgeti, J.-F. Joanny, P. Silberzan, and N. S. Gov, Physics of active jamming during collective cellular motion in a monolayer, Proceedings of the National Academy of Sciences 112, 15314 (2015).

[56] O. Gendelman, Y. G. Pollack, I. Procaccia, S. Sengupta, and J. Zylberg, What determines the static force chains in stressed granular media?, Physical review letters 116, 078001 (2016).

[57] G. Parisi, Y. G. Pollack, I. Procaccia, C. Rainone, and M. Singh, Robustness of mean field theory for hard sphere models, Physical Review E 97, 063003 (2018).

[58] M. Delarue, J. Hartung, C. Schreck, P. Gniewek, L. Hu, and S. Herminghaus, Self-driven jamming in growing microbial populations, Nature Physics 12 (2016).

[59] D. Drasdo, S. Hoehme, and M. Block, On the role of physics in the growth and pattern formation of multi-cellular systems: What can we learn from individual-cell based models?, Journal of Statistical Physics 128, 287 (2007).

[60] S. Turner and J. A. Sherratt, Intercellular Adhesion and Cancer Invasion: A Discrete Simu-lation Using the Extended Potts Model, Journal of Theoretical Biology 216, 85 (2002).

[61] Y. Jiang, J. Pjesivac-Grbovic, C. Cantrell, and J. P. Freyer, A multiscale model for avascular tumor growth, Biophysical journal 89, 3884 (2005).

[62] B. M. Rubenstein and L. J. Kaufman, The role of extracellular matrix in glioma invasion: a cellular potts model approach, Biophysical journal 95, 5661 (2008).

[63] J. F. Li and J. Lowengrub, The effects of cell compressibility, motility and contact inhibition on the growth of tumor cell clusters using the Cellular Potts Model, Journal of Theoretical Biology 343, 79 (2014).

[64] T. J. Newman, Modeling multicellular structures using the subcellular element model, in Single-Cell-Based Models in Biology and Medicine, edited by A. R. A. Anderson, M. A. J. Chaplain, and K. A. Rejniak (Birkhäuser Basel, Basel, 2007) pp. 221–239.

[65] D. A. Matoz-Fernandez, K. Martens, R. Sknepnek, J. L. Barrat, and S. Henkes, Cell division and death inhibit glassy behaviour of confluent tissues, Soft Matter 13, 3205 (2016).

[66] E. Stott, N. Britton, J. Glazier, and M. Zajac, Stochastic simulation of benign avascular tumour growth using the potts model, Mathematical and Computer Modelling 30, 183 (1999).

[67] D. A. Kessler and S. Burov, Effective potential for cellular size control, arXiv preprint arXiv:1701.01725 (2017).

[68] N. Mosheiff, B. M. Martins, S. Pearl-Mizrahi, A. Grünberger, S. Helfrich, I. Mihalcescu, D. Kohlheyer, J. C. Locke, L. Glass, and N. Q. Balaban, Inheritance of Cell-Cycle Duration in the Presence of Periodic Forcing, Phys. Rev. X 8, 21035 (2018).

[69] J. Lin and A. Amir, From single-cell variability to population growth, Physical Review E 101, 012401 (2020).

[70] Z. You, D. J. G. Pearce, A. Sengupta, and L. Giomi, Mono- to multilayer transition in growing bacterial colonies, Phys. Rev. Lett. 123, 178001 (2019).

[71] Note that the cell-sensed pressure is the quantity that determines division rate and on which therefore the above expansion is based, whereas Fig. 4B describes the Irving-Kirkwood pressure. However, we also measured the average cell-sensed pressure in the system and found that the dependence on *E* has identical shape, while systematically being larger by ~ 1% compared to the Irving–Kirkwood pressure in Fig. 4B. Consistently, it is also larger than the zeroth-order approximation of Eq. (F1) for all *E* and our fluctuation-related conclusions in this section remain valid for either pressure definition.

